# Drift drives phenotypic evolution in a rapid island radiation

**DOI:** 10.64898/2026.06.09.731170

**Authors:** Jenna M. McCullough, Chad M. Eliason, Allison J. Shultz, Stepfanie M. Aguillón, David J.X. Tan, Fernando Machado-Stredel, Shannon J. Hackett, Corinne E. Myers, Michael J. Andersen

## Abstract

Understanding the processes that generate phenotypic diversity is central to explaining how new species form^1,2^. Evolutionary theory predicts that rapid evolution of signaling traits, such as feather coloration, can promote speciation^3,4^ but empirical support is inconsistent^5,6^. Phenotypic divergence of such traits is expected during speciation^4^, but these microevolutionary dynamics are rarely examined at macroevolutionary scales or linked to underlying population demography. Here, we leverage complete taxon sampling across an iconic insular bird radiation that helped shape early theories of allopatric speciation. We integrate whole-genome data with a comprehensive, fine-scale dataset of whole-body plumage coloration to directly test whether signaling trait evolution covaries with lineage diversification and to disentangle the roles of selection and drift. We find that lineages with faster rates of color evolution diversify more rapidly. Strikingly, rates of color evolution accelerate as genomic diversity declines, providing direct evidence that genetic drift—rather than strong sexual or ecological selection—can drive rapid phenotypic change in small, isolated insular populations. Together, these results provide compelling evidence that neutral demographic processes can accelerate the evolution of sexual signals and play a central role in generating phenotypic diversity during island radiations.

## Main

The tempo of phenotypic evolution can influence how quickly lineages diversify^1,7^. Rapid evolution of ornamental and signaling traits, such as the colors of anole dewlaps^8,9^, frog skin^10–14^, African cichlid scales^15–17^, or bird feathers^18^, can influence species’ survival and reproduction. In particular, sexual signaling traits offer a powerful lens to examine the relationship between speciation and phenotypic evolution because of their vital role in establishing prezygotic reproductive barriers^17,19–28^. Theory^4^ predicts that, prior to the completion of speciation, phenotypic differences will accrue between diverging lineages reflecting divergent selection on mate recognition between incipient species without requiring measurable shifts in ecological niches^29^. Therefore, higher rates of lineage diversification are expected in conjunction with higher rates of change for signaling traits.

Feather coloration has played a central role in the development of influential speciation theories^3,4^ and is an ideal trait to study links between microevolutionary trait change and macroevolutionary patterns of lineage diversification. Plumage color is tightly integrated with avian ecology, influencing camouflage, thermoregulation, flight performance, and communication^18,30^. Changes in color or pattern can alter social signaling and species recognition^31–33^. Despite strong theoretical expectations, empirical links between rates of color evolution and lineage diversification are sparse and inconsistent^5,6,20,34–36^, reflecting a potential mismatch in evolutionary scale. Sexual signal divergence is predicted to arise early during lineage divergence, yet most comparative analyses quantify phenotypic differences only among species-level taxa, overlooking trait variation that accumulates within geographically structured populations. Resolving this mismatch requires densely sampled datasets that capture divergence both within and among lineages across the speciation continuum.

Three broad hypotheses addressing adaptive or neutral processes could explain microevolutionary plumage divergence during speciation. First, sexual selection^3,29,37^ may accelerate color divergence if shifts in signaling traits alter mate recognition cues, generating prezygotic isolation among diverging populations. Second, selection for species recognition in competitive or ecological contexts may similarly promote divergence as lineages partition color space^38^. In these adaptive scenarios, selection is the primary driver of phenotypic trait evolution. Alternatively, neutral evolutionary processes like genetic drift may contribute to rapid plumage color evolution, particularly in small populations^39–42^. In this case, color divergence would arise through random fixation of variants when effective population sizes are small. Although theory^43–46^ predicts population demography can shape evolutionary trajectories, such predictions have rarely been tested by directly linking population genomic estimates of genetic diversity to phenotypic evolutionary rates across species in natural systems. Because island radiations are subject to founder events and limited gene flow^39–41^, they are a powerful system in which to test whether phenotypic evolution is shaped by adaptive processes or a consequence of demographic history.

Here, we combine comprehensive UV-vis spectrophotometry of plumage color with whole-genome sequencing across the complete taxonomic diversity of *Todiramphus* kingfishers, an insular Indo-Pacific clade whose geographic variation has inspired evolutionary theory^43,47,48^. We quantified whole-body plumage color across 15 patches in 1,045 specimens spanning all 29 species and 96 phenotypically distinct taxa, including extensive sampling of geographically structured populations within species (Figs. 1a, S1). By capturing trait variation across both shallow and deeper levels of lineage divergence, this dataset enables a direct test of how microevolutionary processes scale to macroevolutionary patterns. By integrating phenotypic and genomic data across the complete phylogeny, we test whether rates of signaling trait evolution covary with lineage diversification and whether these dynamics are driven by selection or drift. Our results demonstrate genetic drift as a key driver of rapid phenotypic divergence in this island radiation.

**Fig 1.**
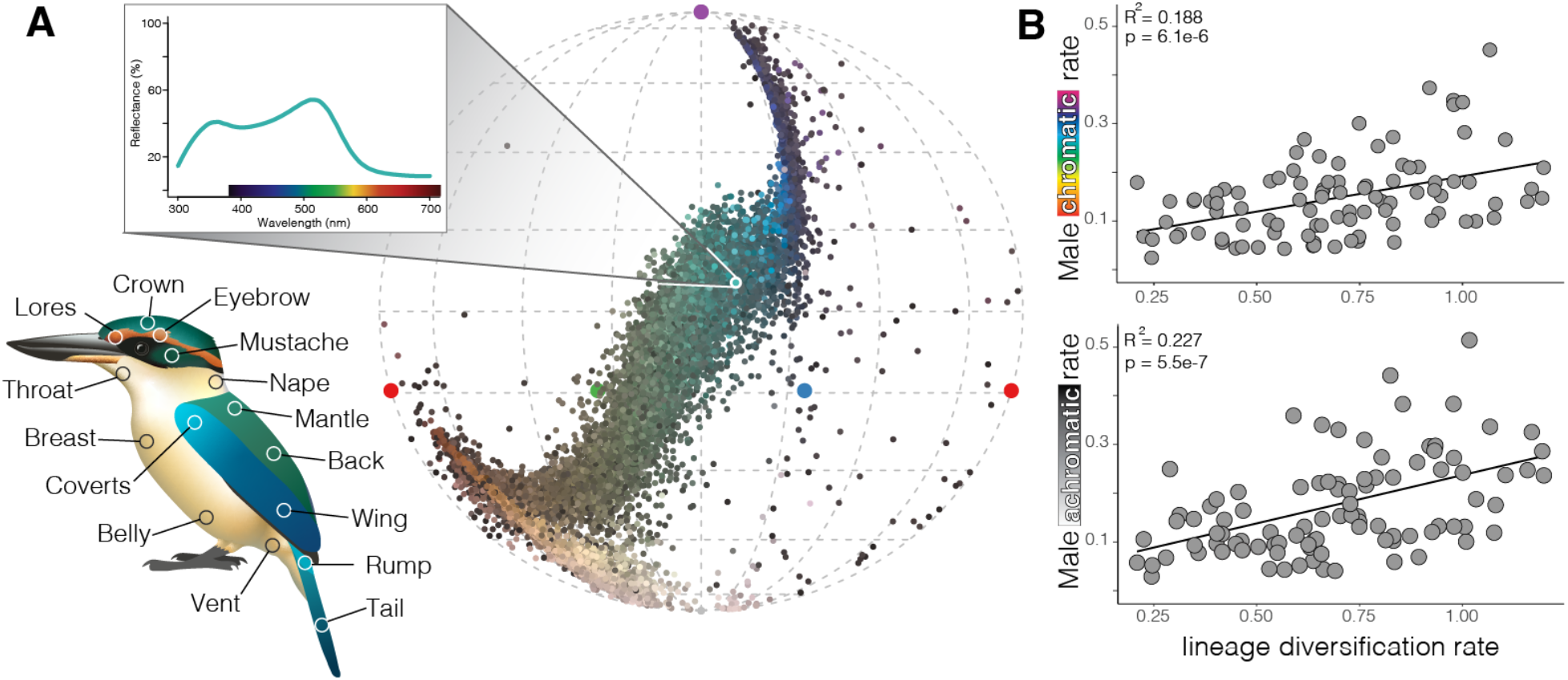
Comprehensive quantification of plumage colors reveals faster phenotypic evolution in rapidly diversifying kingfisher lineages. **a** Two-dimensional projection of four-dimensional tetrahedral color space showing all 15,503 averaged per-patch color measurements from 15 body patches of 1,045 specimens representing 29 species and 96 phenotypically distinct taxa in the genus *Todiramphus* (45,909 total measurements). Each point represents three averaged spectral reflectance measurements (one scan is denoted in the inset graph). Four points (red, blue, green, purple) at the exterior of the circle denote the four types of color cones present in the UV-sensitive visual system. The 15 measured plumage patches are illustrated on a male Pacific Kingfisher (*T. sacer brachyurus*). **b**, Plumage color evolutionary rates increase with lineage diversification: chromatic and achromatic evolutionary rates are positively correlated with diversification rate estimates. Male results are shown here. See Figs. S5–6 for similar results with females and alternative methods of diversification rate estimation.

### Diversification and phenotypic evolutionary rates are positively correlated

Across *Todiramphus* kingfishers, the tempo of plumage color evolution tracks lineage diversification. Lineages exhibiting higher rates of plumage color evolution, measured as either chromatic (hue and saturation) or achromatic (brightness) color, consistently showed higher lineage diversification rates (Fig. 1b). This positive relationship was robust across sexes, axes of color, and alternative methods for estimating lineage diversification rates (Table S1, Figs. S2–6). There was substantial variation in the tempo of both axes of color within species complexes, such Pacific (*T. sacer*, 21 subspecific taxa) and Collared kingfishers (*T. chloris*, 14 taxa; Figs. S3–4), which also exhibit variation in diversification rates (Fig. S2; See Supplementary Material). This within-species heterogeneity is notable given that comparative analyses typically treat species as single units and highlights the importance of accounting for variation within lineages when studying patterns of phenotypic evolution. Together, these results demonstrate a strong, positive relationship between the tempo of phenotypic trait evolution and the accumulation of species diversity in this radiation.

### Tempo of phenotypic evolution increases with decreasing genomic diversity

Having established that rapid color evolution tracks lineage diversification, we next asked whether this pattern reflects expectations consistent with selective mechanisms or neutral evolutionary processes. Sexual selection and interspecific interactions (i.e., competition) are frequently proposed as the primary adaptive drivers of plumage divergence^29,37,49^. However, whole-body measures of sexual dichromatism, a common proxy for the strength of sexual selection^3,34,50,51^, showed no association with rates of color evolution for either sex or color measure (Table S2, Figs. 2a, 2d, S7). Similarly, lineages experiencing greater sympatry with other *Todiramphus*, conditions in which selection for species recognition is predicted to drive divergence as lineages partition color space, did not exhibit faster color evolution (Table S3, Fig. 2b, 2e). Contrary to theoretical predictions, sympatry was weakly associated with lower color evolutionary rates, and only when evolutionary history was not explicitly modelled (Figs. S8–9).

**Fig 2.**
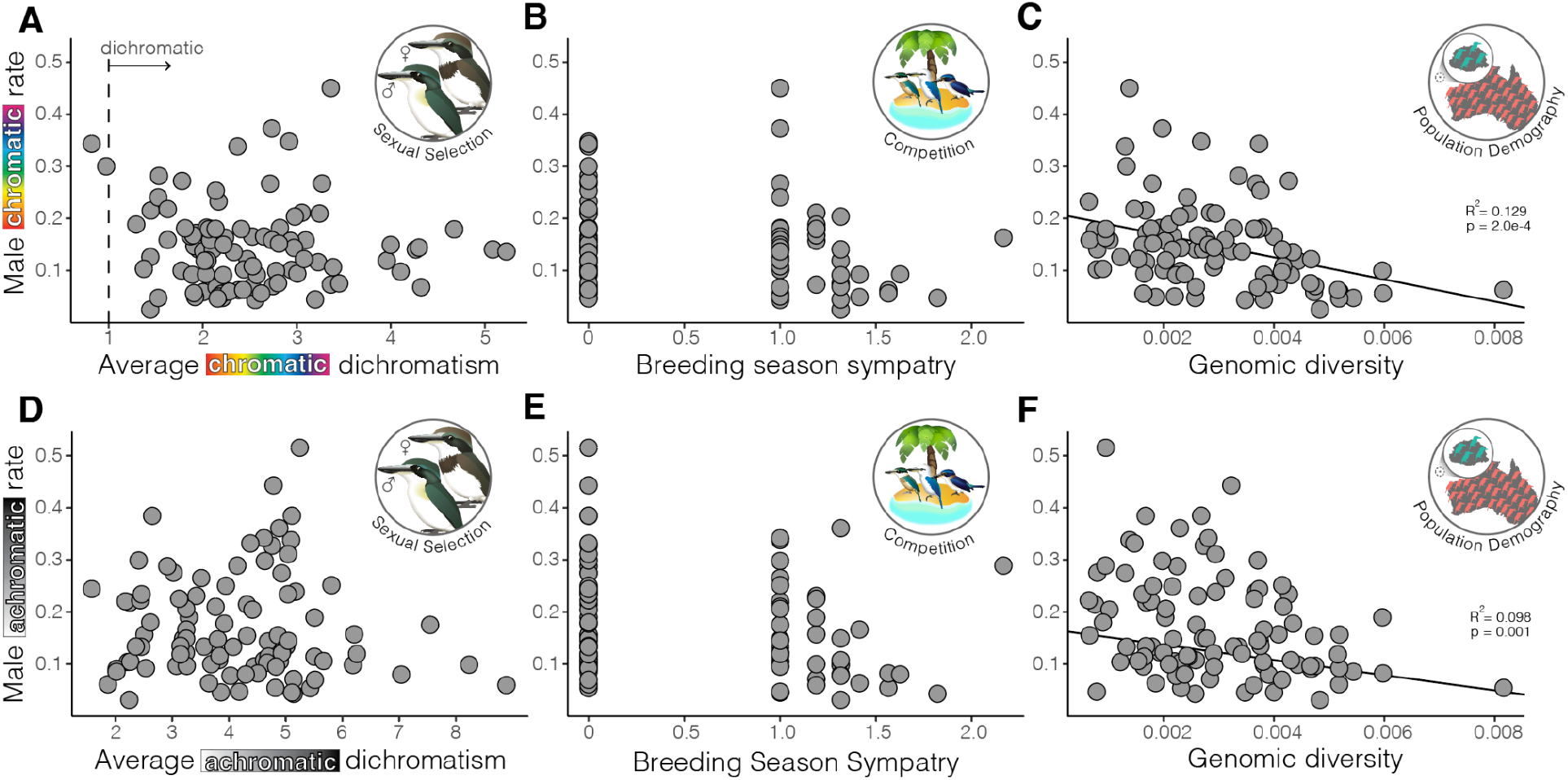
Lower genomic diversity predicts faster plumage color evolution. Male chromatic (hue and saturation; **a**– **c**) and achromatic (brightness; **d**–**f**) evolutionary rates are explained only by heterozygosity (**c**,**f**), a genome-wide measure of genetic diversity and proxy for effective population size. We found no relationship between color evolutionary rates and either chromatic (**a**) or achromatic (**d**) sexual dichromatism, nor with increasing sympatry during the breeding season (**b**,**e**). In panels **a** and **d**, dichromatism values greater than one indicates that males and females are perceptibly different in the avian visual system; note that all taxa are distinguishable for achromatic color. Values of zero in panels B and E indicate taxa that are allopatric during the breeding season. Female evolutionary rates show similar relationships (Figs. S7–11).

Neutral mechanisms like drift are expected to accelerate phenotypic divergence in small populations^44,45^, but the role of drift is rarely demonstrated with empirical data^42^. In lieu of direct census estimates for insular biota, island size is a widely used proxy for population size when testing the effect of drift. However, we found no relationship for island size explaining the tempo of color evolution (Figs. S10–11), consistent with the idea that island area alone may be an incomplete proxy for population size because it does not account for isolation among populations. In stark contrast, genomic diversity was consistently and negatively associated with rates of color evolution (Fig. 2c, 2f). Across 59 models evaluating ecological and genetic predictors, genome-wide estimates of heterozygosity emerged as the strongest and most consistent predictor of chromatic and achromatic evolutionary rates for both sexes (Tables S2–3; Figs. 2c, 2f, S12–14, see Supplementary Material). Lineages with lower genomic diversity consistently exhibited markedly higher rates of chromatic and achromatic evolution (Fig. 3). This pattern is illustrated by widespread and dispersive lineages such as Red-backed (*T. pyrrhopygius*) and Sacred (*T. sanctus*) kingfishers exhibiting higher genomic diversity but low rates of color evolution. Conversely, range-restricted, island-endemic lineages with low genomic diversity show the highest rates of color evolution, such as Pacific (*T. sacer*) and Chattering (*T. tutus*) kingfishers (Figs. S14–15).

**Fig 3.**
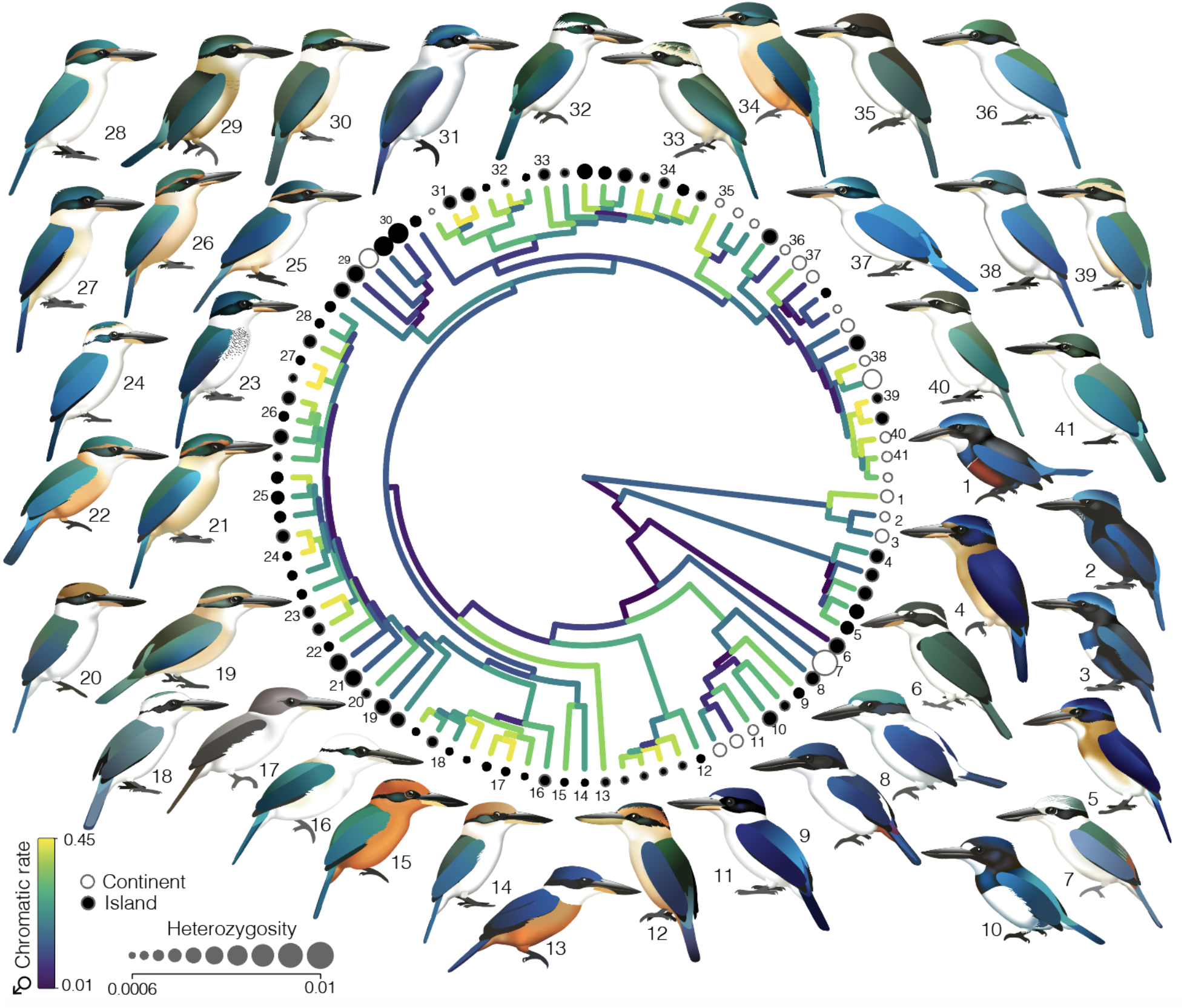
Lineages with lower heterozygosity have higher plumage color evolutionary rates. Maximum clade credibility timetree of *Todiramphus* kingfishers, with branch colors showing the tempo of male chromatic color evolution. Warmer branch colors indicate higher rates of chromatic color evolution. Size of circles at tips reflects genome-wide estimates of heterozygosity. Color of dots indicates whether a species is an island endemic (black) or occurs on continental landmasses and land-bridge islands (white). See Fig. S15 for species names.

### Genetic drift drives rapid phenotypic evolution in an island radiation

Together, these results implicate stochastic drift as a primary driver of rapid plumage color evolution in *Todiramphus*. Reduced genomic diversity in small, isolated populations likely accelerated the fixation of novel color variants in this group. Rather than reflecting relaxed sexual or ecological selection, rapid color divergence in this radiation appears to arise from neutral processes operating early during population divergence. Drift is often treated in macroevolutionary studies as a null expectation to be rejected in favor of adaptive explanations^52–58^. However, evolutionary theory predicts that stochastic processes can accelerate phenotypic divergence, with their effect magnified in small insular populations^59–62^. Our results provide rare macroevolutionary support for this prediction. Despite testing multiple adaptive hypotheses—including sexual selection and interspecific interactions—genomic diversity was the dominant predictor of plumage color evolutionary rates, whereas adaptive factors explained very little to no variation (Supplementary Material, Tables S2–3).

Colonization of islands is widely associated with shifts in phenotypic evolution, reflecting reduced population sizes, founder effects, and increased genetic drift^43,63–66^. The majority of *Todiramphus* kingfishers are endemic to islands (70 of 96 taxa occur only on islands^67,68^). Although island-restricted taxa such as microendemics within Pacific Kingfisher (*T. sacer*) exhibited some of the highest color evolutionary rates and lowest levels of genetic diversity, taxa within continental species also exhibited high evolutionary rates (see *T. chloris*; Figs. 3, S2–4). Studies have shown how drift contributes to within-species divergence of phenotypic traits, such as bat vocalizations^62^, female preference for the colors of poison-dart frog skin^12^, or body size^42^ but this is the first time drift has been identified as the predominant mechanism for phenotypic evolution of a sexual signaling trait on a macroevolutionary scale. Our results show that when demographic history is explicitly considered, rather than inferred using the traditional proxy of island size, stochastic processes can emerge as a driving force in shaping phenotypic evolution in island radiations. This highlights genetic drift as a more potent and underappreciated force shaping phenotypic evolution in small and isolated populations.

We hypothesize that the predominance of non-iridescent structural feather colors in *Todiramphus* kingfishers makes their plumage particularly susceptible to stochastic drift. Avian feather colors are produced through a combination of pigmentary absorption and/or structural light scattering^69,70^, which differ in their developmental basis and evolutionary lability. Pigment-based colors derive from melanins (blacks, browns) and carotenoids (reds, yellows), whereas non-iridescent structural colors (greens, blues, violets) result from the nanoscale arrangement of keratin, air, and melanin that coherently scatter light^71^. Structural colors depend on nanostructure assembly rather than complex biosynthesis and are consequently more evolutionarily labile than pigment-based colors^36,71–77^. Moreover, minute changes in the spacing and shape of this keratin and air matrix produce continuous shifts in hue^78,79^, rendering non-iridescent colors particularly sensitive to drift-driven genetic changes. During feather development, this keratin layer is self-assembled via phase separation of β-keratin^80^, a process that may require fewer coordinated genetic changes than shifts in pigment-based coloration. If structural colors are indeed more susceptible to drift than pigment-based colors, then lineages dominated by melanin- or carotenoid-based plumage should show a weaker relationship between genomic diversity and rates of color evolution, a prediction that can be tested across the many avian radiations in which color production mechanisms vary among close relatives.

## Conclusion

Here, we show that diverse sexual signaling traits can evolve without strong selection and can instead be driven by stochastic processes amplified in small, isolated populations. Low genomic diversity predicts accelerated plumage color evolution across sexes and color axes, identifying drift as a central driver of phenotypic change. The strength of drift may be amplified by the low-constraint nature of structurally produced non-iridescent colors typical of *Todiramphus* kingfishers, where small shifts in keratin nanostructure generate continuous phenotypic variation. By uniting dense taxonomic sampling with genome-wide and phenotypic data, we directly link population-level processes to macroevolutionary patterns. These findings suggest that genetic diversity and mechanism of trait production jointly shape the tempo of phenotypic evolution, and that drift-driven signal divergence may be a common, but underappreciated, precursor to lineage divergence in insular radiations.

## Supplementary Information

**Appendix 1**. Supplementary text (introduction, methods, and results) and figures S1–16.

**Appendix 2**. Supplementary tables

## Author contributions

JMM, CME, and MJA conceived the study. JMM, CME, SJH, CEM, and MJA acquired funding. JMM and CME performed data collection. JMM performed analyses and wrote the original draft with input from all authors. All authors approved the final draft.

## Data & code Availability

Averaged spectral reflectance data is available on the Dryad Digital Repository (Link TBD after acceptance). Code and data to replicate analyses and figures are available on GitHub at https://github.com/JennaMcCullough/Todiramphus_color.

## Acknowledgements

Avian specimens used in this study represent over two centuries (1801 to 2019) of collecting efforts by countless people who have contributed to the growth and use of natural history collections. We specifically acknowledge the curators and collections managers of Australian Museum (AM), American Museum of Natural History (AMNH), The Academy of Natural Sciences of Drexel University (ANSP), Australian National Wildlife Collection (ANWC), Delaware Museum of Natural History (DMNH), the Field Museum (FMNH), University of Kansas Biodiversity Institute and Natural History Museum (KU), Muséum National d’Histoire Naturelle (MHN), Museum of Southwestern Biology (MSB), the Natural History Museum in London (NHM), Queensland Museum (QM), United States National Museum (USNM), University of Washington Burke Museum (UWBM), and Western Australia Museum (WAM). Kristopher Menghi and Nancy Moore assisted with spectral data collection. We thank Ethan F. Gyllenhaal and Devon A. DeRaad for advice and feedback that improved this manuscript. We would like to thank the CIPRES Science Gateway and UNM Center for Advanced Research Computing, both supported in part by the National Science Foundation (NSF), for providing the research computing resources used in this work. This study was funded by NSF (DEB 2112467 to CEM, SJH, and MJA and DBI 2507989 to JMM) and the following sources to JMM: the AMNH Frank Chapman Research Grant, UNM Biology Department Alvin R. and Caroline G. Grove Research and Richard B. Forbes Conservation awards, British Ornithologists’ Union Small Ornithological Research Grant, and the American Ornithological Society Werner and Hildegard Hesse Research award.

## Methods

### Whole-genome sequencing, estimation of timetree and diversification rates

To produce a fully sampled timetree, we re-analysed whole-genome data from recent phylogenomic work on the genus. McCullough et al.^68^ sequenced whole genomes of all described taxonomic diversity within *Todiramphus* at 20X average coverage. Per taxon sampling ranged from one to three museum-vouchered specimens, with genetic data sourced from high-quality tissue or blood samples (40%) or cut from the toepads of dried study skins (60%; Table S5). Datasets used in this study included ultraconserved elements (UCEs^81^) and single nucleotide polymorphisms (SNPs, see ‘Testing Drivers of Color Evolution’). From their 90% complete UCE matrix^82^, we restricted sampling to one representative tip per taxon and produced a 100% complete UCE matrix with Phyluce v.1.7.0^83^. We performed independent Markov chain Monte Carlo (MCMC) runs of 100 million generations for 10 subsets of 50 randomly selected UCE loci^84,85^ (n=100 tips; 500 total unique UCE loci, Table S6) in BEAST v.2.6.6^86^ with CPU acceleration using BEAGLE^87^. We used unlinked site models, incorporated relaxed lognormal clock and birth-death tree models to linked partitions, applied the HKY+G sequence model to each partition, and used the RAxML-inferred topology as a multi-monophyletic constraint prior. We used two island ages^88–91^ and secondary calibrations^84^ for time calibration (see Supplementary Information). We ran each chain twice, resulting in 20 total chains and 2 billion generations. After assessing convergence^92^, we removed 10–35% of trees as burn-in and resampled every 100,000th tree in the posterior distribution to produce 17,120 trees to combine into a single maximum clade credibility (MCC) tree^86^. We implemented two non-model based methods to calculate lineage diversification rate, cladogenetic diversification rate shift model^93–95^ and inverse equal splits (ES) measure^96,97^.

### Colorimetric dataset and analysis of color variation

We measured whole body plumage color for all described taxonomic diversity of *Todiramphus* kingfishers (96 phenotypically distinct taxa, 29 species) from closed-wing museum study skins using UV-vis spectrophotometers (Ocean Optics and Avantes) and pulsed-xenon light sources (following taxonomy from ref^68^; Tables S7–8). For each of the fifteen plumage patches, we measured feather reflectance within the avian-visible spectrum (300–700 nm) at normal incidence in triplicate (n=45 scans per specimen). On average, we measured 10.6 specimens per taxon; for 95% of taxa, we measured at least three specimens for each sex (see Supplementary Information for more sampling details). After including spectral reflectance measurements obtained from recent work on color evolution within kingfishers^98^, our dataset comprised 45,909 total spectra (15,303 after averaging) for 1,045 specimens from 14 natural history collections.

Changes in feather color could be due to chromatic differences, such as changes in the dominant reflected wavelength (hue) or color intensity (saturation), as well as achromatic differences, such as luminance (one of several measures of brightness). Kingfishers are UV-sensitive^99^, therefore, to understand variation for both chromatic and achromatic color within the context of the avian tetrachromatic visual system we modelled quantum catches for each photoreceptor class based on a UV-sensitive visual system using photosensitivity data for the Blue Tit^100^. We projected these stimulation values into a tetrahedral color space (u, s, m, and l channels^101^) using the ‘vismodel’ function from the Pavo v.2.9.0^102,103^ package in R v.4.3.2 ^94^. We converted quantum catches in tetrahedral color space and extracted four datasets: x, y, and z coordinates describing chromatic color and a single value of luminance describing achromatic color of each patch for both sexes. We used the RRphylo v.2.8.0^104^ R package to estimate sex-specific chromatic and achromatic color evolutionary rates across each branch of the *Todiramphus* phylogeny and fitted linear models to test whether diversification rate predicted higher rates of plumage color evolution.

### Testing drivers of color evolution

We used both linear and phylogenetic linear models, implemented with the phylolm v.2.6.2^105^ R package to test three hypotheses to explain rapid color evolution in *Todiramphus* kingfishers: sexual selection, interspecific interactions, and population demography (see Supplementary Methods). First, increased strength of sexual selection could contribute to greater sexual dichromatism, thus explaining higher rates of plumage color evolution because males or females are evolving to become more dichromatic. To assess the degree of sexual dichromatism, we used the Pavo ‘coldist’ function to calculate the perceptually-uniform Euclidean color distances within tetrahedral color space^106^ between males and females for each patch and axis of color. Values larger than one indicate that males and females are differentiable within the avian visual system for a particular patch. We averaged across the per patch indices to produce whole body indices of achromatic or chromatic sexual dichromatism. Second, sympatric taxa may experience selection for more distinctive coloration to improve signal efficacy for mate recognition or interspecific competition. We estimated sympatry by counting the number of *Todiramphus* taxa co-occurring on the same island or in the same region during the breeding or non-breeding seasons based on species distribution maps^107^, taxonomic records^67,108^, and data from regional bird guides^109–121^ (Table S9). We applied a fourth-root transformation prior to fitting linear models due to the highly skewed distribution of sympatry values. Finally, color evolution might occur faster in smaller populations due to neutral, non-adaptive processes. In lieu of direct census counts, we used two proxies for population size: island area and genome-wide estimates of heterozygosity (Pi). To ensure island size reflected isolation during times of low sea levels (no connectivity), we limited analysis to taxa that occur on islands that remained separated from continental landmasses during the Last Glacial Maximum (Table S5). Based on the shapefile of *Todiramphus* taxon distribution maps (Fig. S1), we used R packages sf^122^ v.1.0-24 and terra^123^ v.1.8-93 to extract island polygons and calculate total area (for taxa that inhabit multiple islands) and mean island area. We then log-transformed both metrics prior to analysis. However, island size alone does not account for a population’s degree of isolation. To take genomic diversity into account, we used genome-wide heterozygosity values from recent phylogenomic work^68^, which estimated mean Pi across 50 kb windows with pixy v.1.2.10^124^.

