## Supplemental Text and Figures for "Drift drives phenotypic evolution in a rapid island radiation"

**<u>Appendix 1. Supplementary Material for:</u> Drift drives phenotypic evolution in a rapid island radiation**

**Table of Contents:**

Supplementary Background
Supplementary Methods
Supplementary Results
Supplementary Table Captions (See Appendix 2 for Supplementary Tables)
Supplementary Figures 1–16

### Supplementary Background

*Description of* Todiramphus *Taxonomic and Phenotypic Diversity*

*Todiramphus* comprises 29 species according to current taxonomy and recent genomic work[67,68]. The 96 subspecies or monotypic species (hereafter taxa; Fig. S1) are variably ascribed from the monotypic Pohnpei Kingfisher *T. reichenbachii* (endemic to Pohnpei Island, Micronesia) to the highly polytypic Pacific Kingfisher *T. sacer*, with 23 phenotypically distinct and allopatric subspecies ranging from the Solomon Islands to Samoa[67,68] (Fig. S1). *Todiramphus* kingfishers share similar color patterning, such as white or buffy underparts with dark-blue to pale-green upperparts, but closely related taxa within species complexes can look drastically different (e.g., *T. sacer* in Fig. S15). Many taxa within the *T. sacer* complex have white underparts, whereas the males of *T. sacer ornatus* (Nendo Island, Solomon Islands) and *T. sacer tannensis* (Tanna Island, Vanuatu) are distinctively buffy[107] (Fig. 3). Even within a single subspecies of Beach Kingfisher (*T. s. admiralitatis*), which is endemic to Manus Island within the Bismarck Archipelago, there are two color morphs for crown plumage: white (matching the nominate and widespread subspecies) or greenish-blue (matching the common crown color of the genus[109,126], Fig. S2). Some species are strongly sexually dichromatic, such as Guam (*T. cinnamominus*) and Somber (*T. funebris*) kingfishers, whereas other polytypic species comprise a single taxon that is sexually dichromatic, such as taxa within Tahiti Kingfisher (*T. veneratus veneratus*; Fig. S7) and Blue-Black Kingfisher (*T. nigrocyaneus nigrocyaneus*). As a genus, *Todiramphus* kingfishers are not considered to be strongly dichromatic with the most common description of females as being similar in plumage to the males but with slightly drabber green hues on their upperparts[127,128].

### Supplementary Methods

*Phylogeny Time Calibration and Lineage Diversification*

We used a combination of island ages and secondary calibrations from previous higher-level work on kingfishers and their relatives. We used two secondary calibrations from McCullough et al.[129], which produced a completely sampled species timetree of the order Coraciiformes based on three fossils and two secondary calibrations from a higher-level timetree of all birds[130]. For our analyses, we used two calibrations with normal distributions on the node subtending Blue-capped Kingfisher (*Actenoides hombroni*) with a mean date of 17.28 Ma (95% confidence interval=19.7–14.8 Ma, sigma=1.5) and the node subtending Gray-headed Kingfisher (*Halcyon leucocephala*) and crown-group *Todiramphus* with a mean date of 10.33 Ma (CI=8.37–12.3 Ma, sigma=1.0). We applied two island age calibrations for two pairs of Polynesian taxa. Specifically, we placed a uniform distribution with an upper bound of 1.53 Ma on the split of two Pacific Kingfisher (*T. sacer*) taxa that are endemic to adjacent islands in American Samoa, *pealei* (single island endemic of Tutuila) and *manuae* (endemic to islands of Manu'a). Tutuila Island is part of the Samoa Hot Spot and is estimated to have been subaerial by 1.53–1.0 Ma[88,89]. The second island age we used was the age for Tahiti, which dates the split between the two taxa of Tahiti Kingfisher (*T. veneratus*) that are endemic to the Windward Islands within the Society Islands, French Polynesia. *Todiramphus v. veneratus* is endemic to Tahiti (1.2–0.6 Ma) and *T. v. youngi* is endemic to the adjacent and older island of Moorea (2.0–1.5 Ma[88,90,91]). We placed a uniform distribution on this node with an upper bound of 1.2 Ma. We ran each of the 10 UCE subsets twice, resulting in 20 total chains and 2 billion generations. We visualized posterior estimates and assessed convergence in Tracer v.1.7.1[92]. We discarded 10–35% of trees as burn-in before combining the two independent runs for all ten subsets in TreeAnnotator v.2.6.7[86]. We resampled every 100,000th tree in the posterior distribution to produce 17,120 trees that we combined into a single maximum clade credibility (MCC) tree.

With a completely sampled phylogeny of all *Todiramphus* taxa, we estimated the lineage diversification rate across the genus. Though studies that estimate diversification rates are largely conducted at the species level, our approach to estimate diversification rates here is justified because we have complete sampling of all taxonomic diversity in the genus (as opposed to species-level sampling for some species complexes and subspecific sampling in others, which would introduce bias). In light of the debate[131–133] on the best methods to investigate macroevolutionary trends, we implemented two non-model-based methods to estimate lineage diversification rate in order to consider how they impact inferences of plumage color evolution. First, we used the cladogenetic diversification rate shift model (CLaDS[93]), implemented in the R[94] package RPANDA v2.2[95]. CLaDS is a Bayesian program that estimates branch-specific speciation rates and uses an autocorrelated clock model to estimate small, frequent shifts in speciation rate across phylogenies. This method has been highlighted to perform better than models inferring single large shifts in speciation rate within birds[134]. Second, we compared these values with the lineage diversification rates calculated by the inverse equal splits (ES) measure[96,97], estimated in R.

*Colorimetric Dataset Sampling*

We used two spectrophotometers, either an Avantes AvaSpec or Ocean Optics with pulsed xenon light sources (AvaLight-XE or PX-2, respectively) to measure reflectance within the bird-visible spectrum (300–700 nm) from closed-wing museum study skins (Table S7). We took triplicate measurements at normal incidence of 15 plumage patches, resulting in 42 measurements for each specimen. We collected colorimetric datasets for both sexes because recent work has highlighted decoupled rates of plumage evolution between male and females of the same species[6] and some species complexes like the Rufous-lored Kingfisher (*T. winchelli*, Philippines) display more plumage color differences for females rather than males.

We measured plumage color for all described diversity of *Todiramphus* kingfishers. On average, we measured 10.6 individuals per taxon (Table S3). For 95% of taxa, we measured at least six individuals total for each taxon and three individuals for each sex (Tables S2–3). Because of the scarcity of specimens in museum collections, there were four taxa for which we were not able to measure six individuals: two taxa of the Cinnamon-banded Kingfisher (*T. australasia odites* [n=3] and *tringorum* [n=5]), one of the Pacific Kingfisher (*T. sacer vicina* [n=4]). Only partial skins of *T. chloris kalbaensis* exist, which limited whole body colorimetric measurements (this taxon was removed for some downstream analyses that required full body measurements). McCullough et al.[68] found two unnamed, single island endemic populations of *T. sacer* as distinct, either endemic to Tucopia Island, Solomon Islands (*T. sacer tucopia*) or Aneityum Island, Vanuatu (*T. sacer aneityum*). When originally described in 1931, these populations were considered phenotypically distinct based on plumage and morphometric differences but were not named as their own respective subspecies[47]. Because genetic samples from these populations highlight that they warrant their own taxonomic rank, they are included here as *T. sacer tucopia* and *T. sacer aneityum* (McCullough et al. *in prep*). We measured all specimens representing these two unnamed populations of *T. sacer* available in any natural history collection: two of each sex for *T. sacer tucopia* and four male *T. sacer aneityum*. Additionally, we sampled multiple individuals for two microendemic Beach Kingfisher taxa that are unique in the genus by exhibiting two head-color morphotypes, white or blue-green (*T.*

*saurophagus admiralitatis* and *T. s. anachoreta*). We measured more individuals for widespread taxa like the migratory Sacred Kingfisher (*T. sanctus*; n=36) and Collared Kingfisher (*T. chloris humii*, Southeast Asia; n=37) or that live on many islands, such as taxa within Collared Kingfisher (*T. chloris collaris*, Philippines, n=49; *T. chloris chloris*, Wallacea, n=35) and Pacific Kingfisher (*T. sacer vitiensis,* Fiji, n=32; *T. sacer juliae*, Vanuatu, n=48; see Table S8). Because McCullough et al. [68] found Talaud Kingfisher (*T. enigma*) embedded within the *T. chloris* species complex, we treat it here as a taxon of *T. chloris* in species-level taxonomy (such as when plotting dot colors as in Fig. S5). After supplementing our own measures with spectral reflectance measurements from other recent studies on color evolution in kingfishers[98], our dataset comprised 45,909 total spectra (15,303 after averaging) for 1,045 specimens (Table S7).

*Assessing Variation in Color*

With the v.2.9.0 Pavo[102,103] R package, we first averaged the triplicate per-patch reflectance spectra for each of the 15 patches per specimen. Kingfishers are UV-sensitive[99], therefore we modelled quantum catches for each photoreceptor class using photosensitivity data for the Blue Tit[100]. With the 'vismodel' and 'colspace' functions in Pavo, we projected these stimulation values into a tetrahedral colorspace (u, s, m, and l channels[101]). From these data, we extracted per-patch XYZ tetrahedral colorspace coordinates and luminance values for all individuals. The XYZ coordinates describe chromatic (hue and saturation) color, whereas luminance (brightness) describes the achromatic color. We treated these two metrics of color separately because they could be under varying selective pressures[135,136] and to allow for direct comparisons with previous work on coloration in island birds[98,137]. Treating males and females separately, we averaged per-patch X, Y, and Z coordinates and luminance values for individuals within each taxon. This reflects the average colorspace variables per patch for each taxon, with the exception of two Beach Kingfisher taxa (*T. saurophagus admiralitatis* and *T. s. anachoreta*). These taxa are unique in the genus as they each have two different plumage morphotypes for head color (white or blueish-green crowns), which likely reflects a derived character. Therefore, we chose to only analyse data from individuals with blueish-green crowns for these two taxa. Overall, our final four datasets for macroevolutionary analyses are as follows: chromatic (averaged per-patch X, Y, and Z coordinates; 45 trait columns) and achromatic (averaged per-patch luminance; 15 trait columns) color datasets for males and females.

We estimated lineage-specific rates of trait evolution of chromatic and achromatic plumage colors using the v.2.8.0 RRphylo[104] R package. RRphylo uses phylogenetic ridge regression[138,139] to infer evolutionary rates across each branch of a phylogeny. This allows for phenotypic evolutionary rates to vary at each branch, thus allowing this method to be more sensitive to small, incremental changes in phenotype along individual branches rather than relying on large clade-wide shifts to infer phenotypic evolution[140]. We used RRphylo's 'auto.recognize' function to explore the possibility of single clade rate shifts. Two lineages did not have sufficient spectral data for lores (specifically *T. tristrami bennetti* and *T. lazuli* for both sexes and female *T. chloris davisoni*) because this patch was most likely to be dirty due to specimen preparation techniques. Because some comparative methods do not allow for missing data, we opted to drop the lore patch for this specific analysis in an effort to analyse all *Todiramphus* taxa. We fit linear models in R to test whether diversification rate predicted color evolutionary rate.

*Testing Drivers of Color Evolution*

We tested three hypotheses to explain rapid color evolution in *Todiramphus* kingfishers. For each predictor dataset, compared traditional linear models with phylogenetic linear models[105] in R to evaluate whether explicitly accounting for shared evolutionary history altered inference. Although evolutionary rates are estimated from a phylogeny and therefore inherently incorporate phylogenetic information, phylogenetic linear models provide a conservative test by modelling covariance among lineages. If multiple models were significant ($p < 0.05$), we performed multiple linear regression models and assessed the best-fit with Akaike information criterion (AIC) in R.

Sexual selection is widely regarded as a driver of plumage color evolution[29,37] and its strength is often measured by the degree of sexual dichromatism, in which more pronounced differences between males and females suggest stronger sexual selection than when the sexes appear similar[3,34,50,51]. Differences in sexual dichromatism among diverging populations may lead to prezygotic isolation if shifts in signaling traits lead to mismatched mate recognition. Because sexual dichromatism has been linked with rapid color evolution[29,37,49], more dichromatic lineages may exhibit higher diversification rates at macroevolutionary scales. Therefore, we predicted that lineages with stronger sexual dichromatism would have the highest color evolutionary rates. To calculate the degree of sexual dichromatism, we used the Pavo 'coldist' function to calculate the perceptually-uniform Euclidean color distances with tetrahedral colorspace[106] between males and females for each patch and axis of color. This produces two values: an average dichromatism index (i.e., "just noticeable differences", JNDs) for chromatic and achromatic values for each of the 14 patches. Values larger than one indicate that males and females are differentiable within the avian visual system for a particular patch. We averaged across the per patch indices to produce whole body indices of achromatic or chromatic sexual dichromatism. Because we did not have enough specimens for each sex of *T. chloris kalbaenesis* and *T. s. aneityum,* we dropped them from the dichromatism dataset. Whole body measures of dichromatism for *T. lazuli, T. tristrami bennetti,* and *T. chloris davisoni* are derived from averages from 14 of the 15 patches because these species did not have quality data for the lore patch.

The second hypothesis concerned whether sympatric taxa may experience selection for more distinctive coloration to improve signal efficacy for mate recognition or interspecific competition. With increasing degree of sympatry, taxa would have higher rates of plumage color evolution. We estimated sympatry by counting the number of *Todiramphus* taxa co-occurring on the same island or in the same region, based on species distribution maps[107], taxonomic records[67,108], and data from regional bird guides[109–121]. The degree of sympatric *Todiramphus* varies widely. Sympatry varies seasonally for the three migratory taxa (*T. sanctus sanctus*, *T. macleayii macleayii*, *T. macleayii incinctus*) whereas 40 taxa do not co-occur with another *Todiramphus* because most are single-island endemics. Others can be highly sympatric: the nominate taxon of Beach Kingfisher (*T. s. saurophagus*) overlaps with 22 *Todiramphus* taxa across its range from Halmahera to the Solomon Islands (Table S9, Fig. S1). Similarly, the migratory Sacred Kingfisher (*T. s. sanctus*) breeds in Australia and overlaps with six other taxa but may co-occur with up to 42 taxa during the non-breeding season[107]. Due to this wide variation and the skewed distribution of sympatry values, we applied a fourth-root transformation to the sympatry data prior to hypothesis testing with linear models.

Finally, rapid color evolution could result from neutral, non-adaptive processes such as drift, which has a stronger effect in small populations—a well-documented feature of island endemic species. Although population size can be estimated through direct counts[141] or genomic estimates of effective population size[142], these approaches are either labour-intensive or require high-coverage genomes, which are not available for much of the *Todiramphus* clade. As practical alternatives, we used two different proxies for population size: 1) island area and 2) genome-wide estimates of heterozygosity (Pi) from recent phylogenomic work on the genus[68].

*Island Area*—*Todiramphus* kingfishers inhabit a range of islands, from single island endemics, such as *T. sacer regina* (endemic to Futuna Island of the French Territory of Wallis and Futuna), to hundreds of islands, like *T. tristrami alberti* (found across the western and central Solomon Archipelago from Buka to Guadalcanal). Because not all *Todiramphus* taxa are island-restricted, we limited the island-area proxy to the 75 of 96 taxa that occur exclusively on islands (Table S5, Fig. S1). To ensure these values reflected isolation during times of low sea levels (no connectivity), we further restricted analyses to taxa inhabiting islands that remained separated from continental landmasses during the Last Glacial Maximum (LGM). For instance, *T. chloris palmeri* (endemic to Indonesian Islands of Java and Bali; Fig. S1) was excluded because these islands were connected to Southeast Asia via the Sunda Shelf, whereas *T. chloris azelus*, endemic to Enggano Island, was included because this island remained isolated during the LGM. We edited a shapefile of individual, taxon-specific ranges (similar to Fig. S1) in Adobe Illustrator with information from species distribution maps[107], taxonomic records[67,108], and data from regional bird guides[109–121]. We projected this shapefile to the Equi7 Oceania projection and extracted and calculated the area ($m^2$) of the islands in which each taxon occurs with sf [122] v.1.0-24 and terra[123] v.1.8-93 R packages. We considered the total subaerial area of islands in which a taxon occurs, representing its total range size. Because this could be an overestimation of the average island size for taxa that live on multiple islands, we also calculated the mean island area and log transformed both of these metrics prior to analysis.

*Genomic Diversity*—Heterozygosity has been consistently shown to scale with effective population size, with smaller populations having lower estimates of heterozygosity[143,144]. McCullough et al.[68] estimated genome-wide heterozygosity (Pi) for all *Todiramphus* taxa with pixy 1.2.10[124], averaged across 50 kb windows. They summarized genome-wide Pi as the total number of pairwise differences across all windows (sum of count_diffs) divided by the total number of non-missing pairwise comparisons (sum of count_comparisons) thereby yielding a single genome-wide estimate averaged across all windows. Because DNA sourced from museum samples is known to exhibit biases due to degradation[145], and although pixy accounts for missing data when estimating Pi, we fit linear and phylogenetic linear models on two heterozygosity datasets for each sex: 1) all taxa (n=96) and 2) only taxa whose estimates are derived from modern tissue samples (n=44).

### Supplementary Results

#### *Time Calibration and Lineage Diversification*

Our time-calibrated phylogeny inferred a crown age of *Todiramphus* at 10.89 Ma (95% HPD=8.98–12.85 Ma), with early branching lineages (*T. nigrocyaneus*, *T. winchelli*, *T. funebris*, and *T. pyrrhopygius*) diverging prior to the Miocene-Pliocene boundary (Fig. S2). The overwhelming diversity of *Todiramphus* arose within the last five Ma, with substantial differences in species richness between *Todiramphus* clades. For example, a small clade of five species (referred to hereafter as the "*leucopygius* clade", inclusive of *T. albonotatus* and *T. macleayii*) diverged around the Miocene boundary (5.2 Ma; range 3.8–6.8 Ma ) and diversified into five species since its most recent common ancestor (MRCA) 3.3 Ma (2.05–4.74 Ma). At a similar time, we inferred the split between the two most speciose clades of the genus (3.4 Ma; 2.7–5.1 Ma), the Oceanic clade (inclusive of *T. pelewensis* and *T. sacer*) and the Australasian clade (inclusive of *T. sanctus* and *T. chloris*; Fig. S2, clade names follow McCullough et al.[68]). These two clades show a marked increase in diversification rates, with the Oceanic clade accumulating 11 species (34 taxa) since its MRCA at 3.17 Ma (2.19–4.2 Ma) and the Australasian clade diversifying into eight species (39 taxa) since its MRCA at 3.14 Ma (2.17–4.14 Ma). Within the Oceanic clade, a smaller clade of microendemic taxa in east Polynesia (inclusive of Marquesan Kingfisher *T. godeffroyi* and Chattering Kingfisher *T. tutus*, referred to as the "East Polynesian clade") are some of the most recently diversifying species-level taxa in the dataset, having diversified into five species (8 taxa) since 1.42 Ma (0.86–2.05 Ma).

Crown ages for polytypic species complexes are variable but within the last 3 Ma. The two most recent species-level splits are related to island endemic species, either in East Polynesia between Niau Kingfisher (*T. gertrudae*) and *T. tutus* (0.86 Ma, range 0.46–1.29 Ma) as well as widespread Beach Kingfisher (*T. saurophagus*) and island endemic Mariana Kingfisher (*T. albicilla*; 1.03 Ma, range 0.53–1.63 Ma) in Micronesia. These divergence dates are substantially younger than the crown ages for some of the other superspecies complexes, for which the oldest crown age is 2.35 Ma for *T. sanctus* (five total taxa, crown age range 1.46–3.29 Ma). The two most polytypic species complexes, Pacific Kingfisher (*T. sacer*) and Collared Kingfisher (*T. chloris*) have similar crown ages (1.69 and 1.75 Ma, respectively) and both of these groups showed some of the fastest diversification rates estimated by both CLaDs and ES diversification rate metrics (Fig. S2, 16). Overall, diversification rates estimated from our time-calibrated phylogeny were largely concordant, though the ES measure inferred higher rates of lineage diversification than CLaDS (Fig. S16).

*Positive Link between Phenotypic Evolution and Diversification*

The tempo of phenotypic evolution and diversification varied across *Todiramphus*, with lineages showing the highest rates of color evolution also exhibiting the highest lineage diversification rates (Figs. S5–6). We consistently found a statistically supported and positive relationship for diversification rate and color evolution for both sexes, as well as for types of color (chromatic and achromatic) and methodology for estimation of diversification rates (p-value range=0.04–$7.8e^{-9}$; Table S1). We found substantial variation in the rate of color evolution between males and females across *Todiramphus* kingfishers (Figs. S3–4), with no support of a single clade rate shift for any analysis.

*Testing Drivers of Color Evolution*

Across 59 models, genomic heterozygosity emerged as the most consistent predictor of plumage color evolutionary rates across both color axes and sexes (Tables S2–3). For chromatic color evolution, heterozygosity was the sole predictor supported for both males and females under both modelling frameworks (linear and phylogenetic linear models), indicating that inference is robust to whether phylogeny is modelled explicitly.

For achromatic color evolution, phylogenetic linear models again supported heterozygosity as the only predictor, whereas traditional linear models identified weak additional effects of sympatry (Tables S2–3). This sympatry effect explained little additional variance and was opposite to adaptive predictions, with higher achromatic evolutionary rates in allopatric and isolated lineages (Table S3, Figs. S8–9). In linear models of male achromatic evolution, heterozygosity alone explained a substantial proportion of variance (Adj. $R^2$=0.11), while sympatry in either the breeding or non-breeding season was individually significant but explained little additional variance (Adj. $R^2 \leq 0.05$). A multiple linear regression including heterozygosity and breeding-season sympatry yielded the best-supported model, though heterozygosity remained the dominant predictor (Table S4). For females, heterozygosity was again the strongest single predictor of achromatic evolutionary rates. Although breeding-season sympatry was individually significant in linear models, it explained comparatively little variance, and in the best-supported multiple regression model only heterozygosity remained statistically significant.

Comparisons of heterozygosity between samples from different sources (i.e., high-quality tissue versus toepad clippings from historical museum study skins) suggested that heterozygosity may be artifactually inflated in toepad-derived samples (Figs. S12–13). Yet, when we excluded taxa whose heterozygosity estimates were based on toepad-sourced data, the negative relationship between heterozygosity and color evolution was still statistically significant (Tables S2–3), confirming that this relationship is not driven by sample source.

Other predictor variables, sexual dichromatism and island size, were not significant across models. All but two taxa, *T. veneratus youngi* (Moorea, Society Islands) and *T. saurophagus anachoreta* (Anchorite Islands, Papua New Guinea), are identifiably dichromatic in the UV visual system for chromatic colors (values larger than 1 in Fig. S7A–B); all *Todiramphus* taxa are dichromatic for achromatic colors. But dichromatism did not show a relationship to rates of color evolution in either axis (Table S7). Similarly, either metric of physical island size—whether defined as the total or mean area of islands on which a single taxon occurs—did not show a relationship to the tempo of color evolution (Figs. S10–11).

### Supplementary Table Captions

*See Appendix 2 for Supplementary tables in Excel format*

**Table S1.** Linear models testing the relationship between chromatic (hue and saturation) and achromatic (brightness) color evolutionary rates and lineage diversification rates in male and female *Todiramphus* kingfishers.

**Table S2.** Summary of single-predictor phylogenetic linear models examining effects of sympatry, sexual dichromatism, island size, and genomic diversity (heterozygosity) on chromatic (hue and saturation) and achromatic (brightness) color evolution rates in male and female *Todiramphus* kingfishers. For each model, we report the model p-value and number of taxa excluded due to missing data (e.g., absence from dichromatism or island datasets, or heterozygosity estimates from toepad-sourced DNA). If significant, we report the adjusted $R^2$ and coefficients. Model terms are as follows: chromatic evolutionary rate (colrate), achromatic evolutionary rate (lumrate), breeding season sympatry (symp_breed), year-round sympatry (symp_all), chromatic dichromatism (AvgDichromCol), achromatic dichromatism (AvgDichromLum), log transformed island total area (island_size_log_totalarea), log transformed island mean area (island_size_log_meanarea).

**Table S3.** Summary of single-predictor linear models examining effects of sympatry, sexual dichromatism, island size, and genomic diversity (heterozygosity) on chromatic (hue and saturation) and achromatic (brightness) color evolution rates in male and female Todiramphus kingfishers. For each model, we report the model p-value and number of taxa excluded due to missing data (e.g., absence from dichromatism or island datasets, or heterozygosity estimates from toepad-sourced DNA). If significant, we report the adjusted $R^2$ and coefficients. Model terms are as follows: chromatic evolutionary rate (colrate), achromatic evolutionary rate (lumrate), breeding season sympatry (symp_breed), year-round sympatry (symp_all), chromatic dichromatism (AvgDichromCol), achromatic dichromatism (AvgDichromLum), log transformed island total area (island_size_log_totalarea), log transformed island mean area (island_size_log_meanarea).

**Table S4.** Summary of single and multiple-predictor linear models evaluating the effects of genetic diversity, sympatry, and sexual dichromatism on sex-specific chromatic (saturation and hue) and achromatic (brightness) evolutionary rates. Shown are adjusted $R^2$, F-statistics, model and term p-values, degrees of freedom, and AIC. See Table S2 for all single-predictor linear models.

**Table S5.** Chromatic (color) and achromatic (luminance) rates of plumage color evolution and dichromatism in *Todiramphus* kingfishers. Genome-wide heterozygosity estimates are from McCullough et al. (2025). Mean and total island area are in meters. Average dichromatism indices are calculated from all 15 patches. For information as to why some taxa have missing data for dichromatism and island data, see Supplementary Material.

**Table S6.** Ten unique subsets of 50 randomly selected UCE loci used for time calibration in BEAST.

**Table S7.** Specimens measured for this study. Collection abbreviations are as follows: AM, Australia Museum; AMNH, American Museum of Natural History; ANSP, Academy of Natural Sciences of Drexel University; ANWC, Australian National Wildlife Collection; DMNH, Delaware Museum of Natural History; FMNH, the Field Museum; KU, University of Kansas Biodiversity Institute and Natural History Museum; MHN, Muséum National d'Histoire Naturelle; MSB, Museum of Southwestern Biology; NHM, the Natural History Museum in London; QM, Queensland Museum; USNM, United States National Museum; UWBM, University of Washington Burke Museum; and WAM, Western Australia Museum. Because of different numbering conventions across collections, some spectral reflectance scans have alternative accession numbers for scans

**Table S8.** Taxon- and sex-specific breakdown of specimens measured for this study.

**Table S9.** Number of sympatric *Todiramphus* taxa during the breeding and non-breeding seasons.

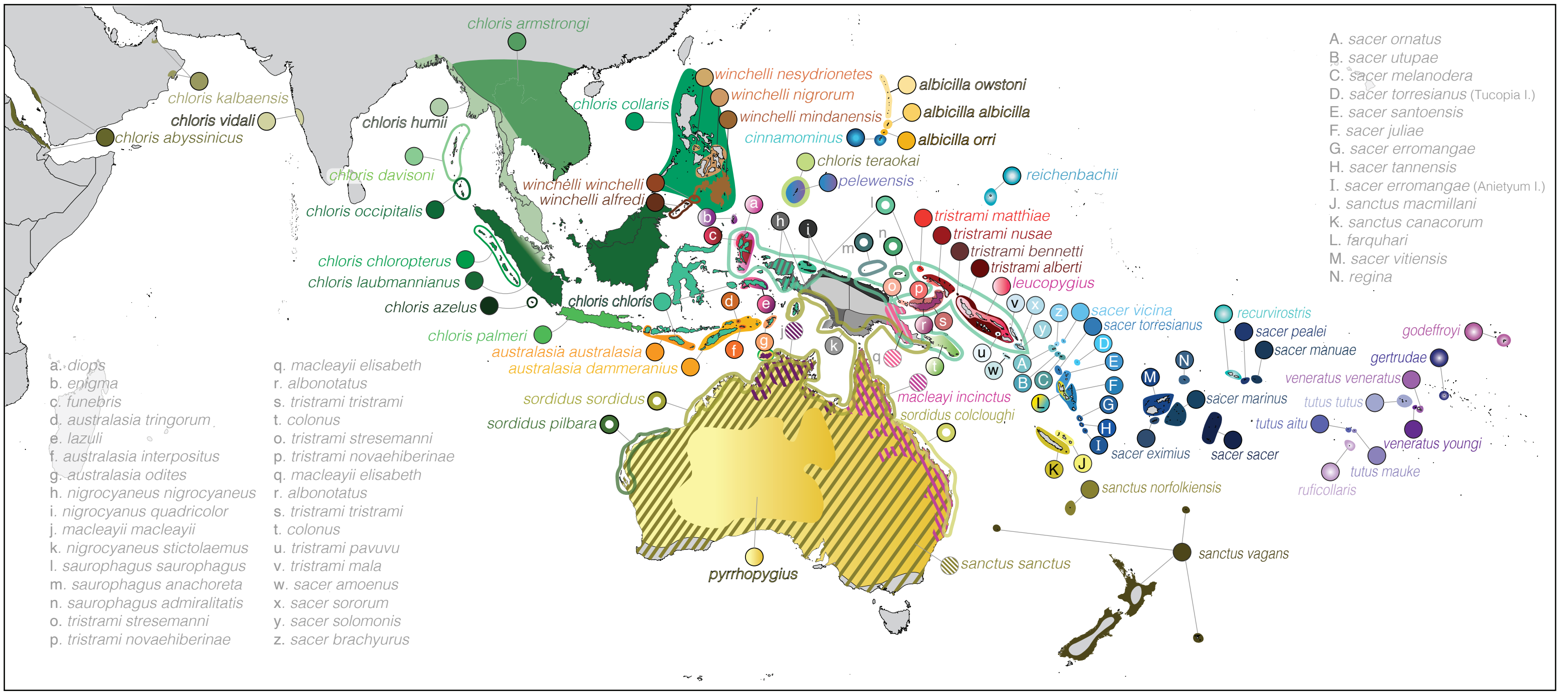


Figure S1. Map of *Todiramphus* kingfishers. Taxon-specific breeding ranges of all 96 extant taxa within the genus, with diagonal lines showing sympatry in some cases. Adapted from McCullough et al. 2025.

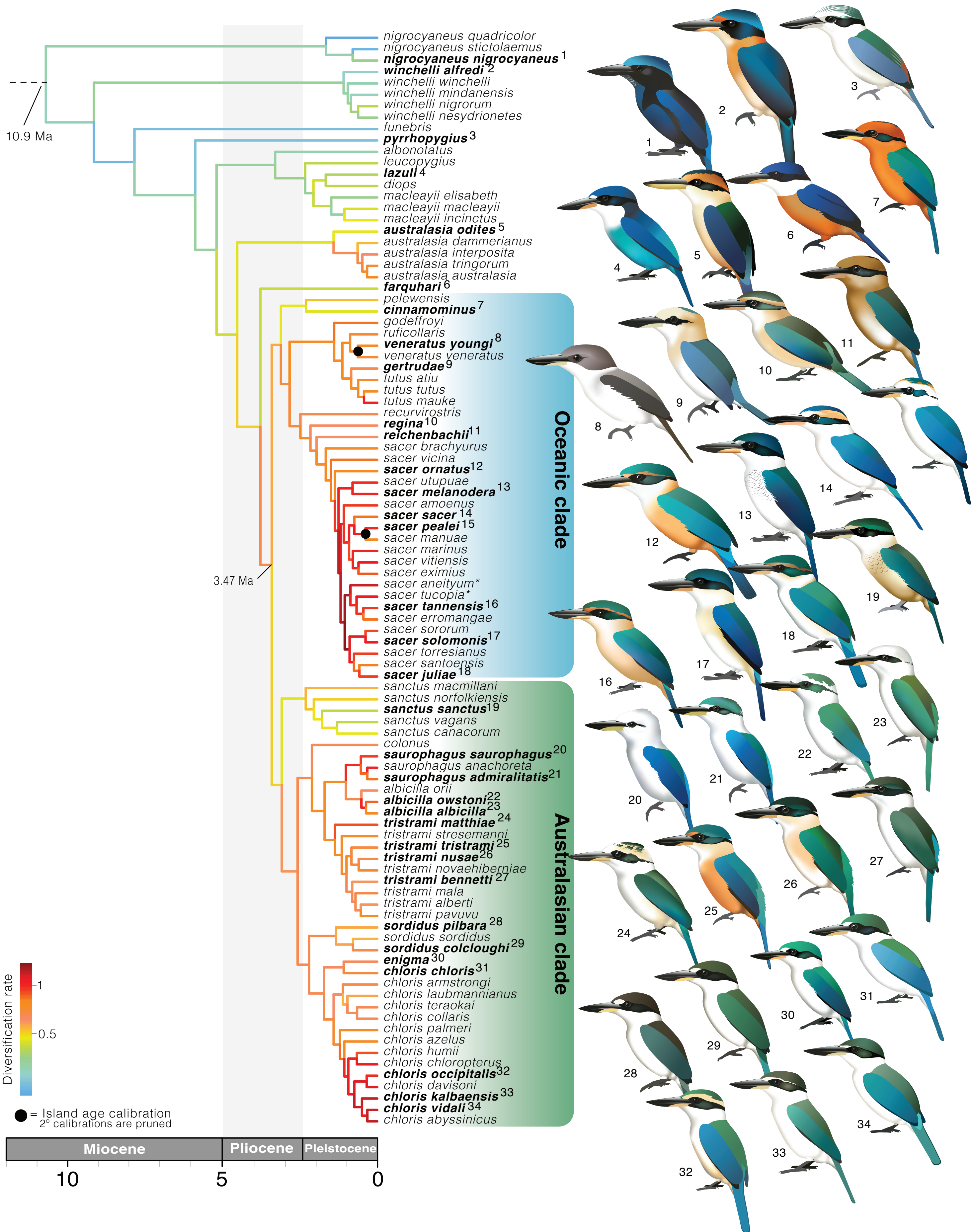


Figure S2. Maximum clade credibility (MCC) tree of 500 ultraconserved loci and fixed RAxML topology produced in BEAST. Placement of secondary calibrations are not shown because they were on outgroup nodes, pruned herein. Calibration points for island ages are shown as black circles. Branch colours correspond to diversification rates estimated with the cladogenetic diversification rate shift (CLaDs) model. Illustrations of representative taxa correspond to numbered tip labels and were created by Jenna McCullough.

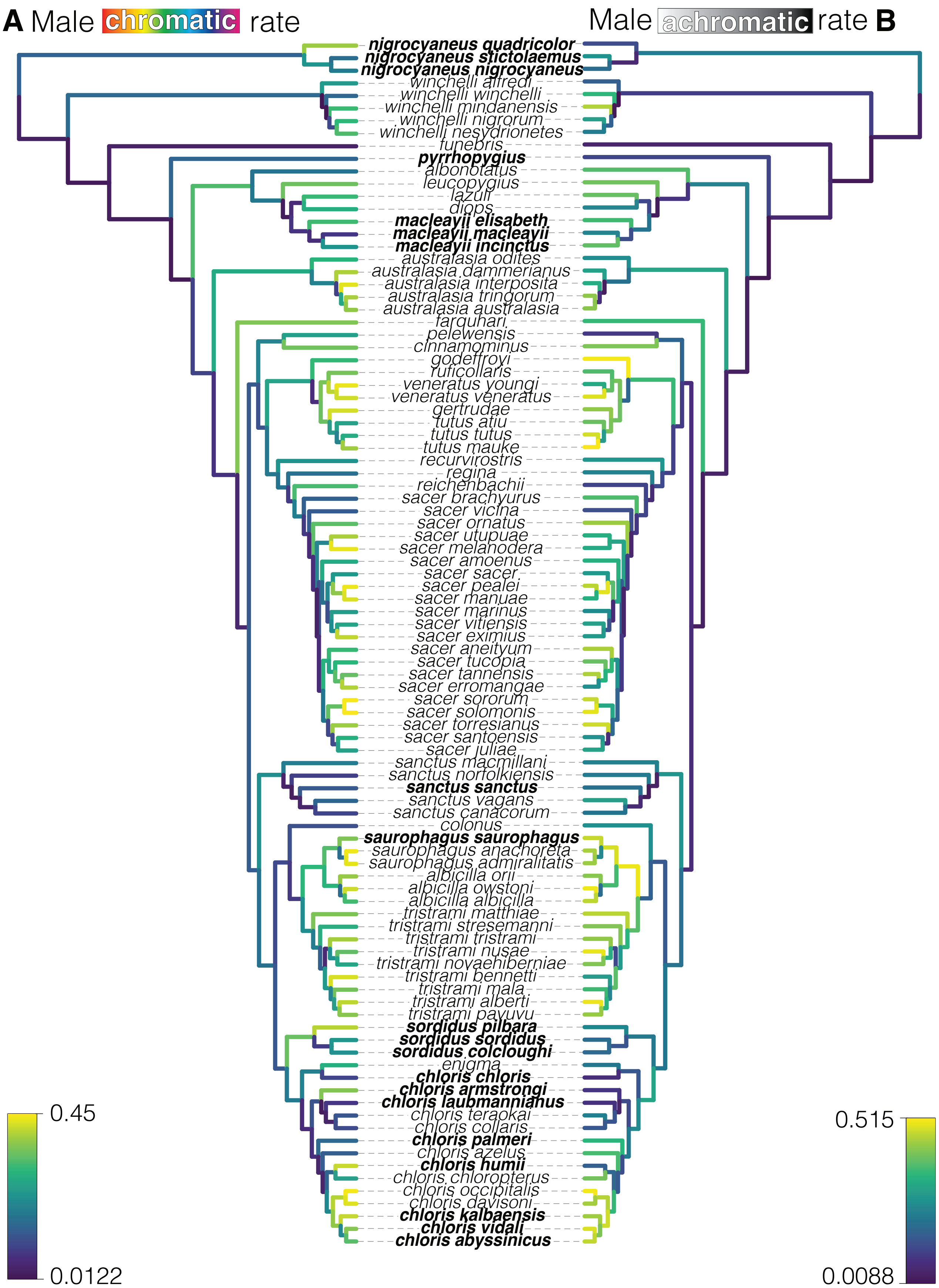


Figure S3. Tempo of male plumage colour evolution for A) chromatic (describing hue and saturation) and B) achromatic (describing brightness) colours across 14 plumage patches. Warmer branch colours indicate higher rates of colour evolution. Bolded names indicate continental taxa.

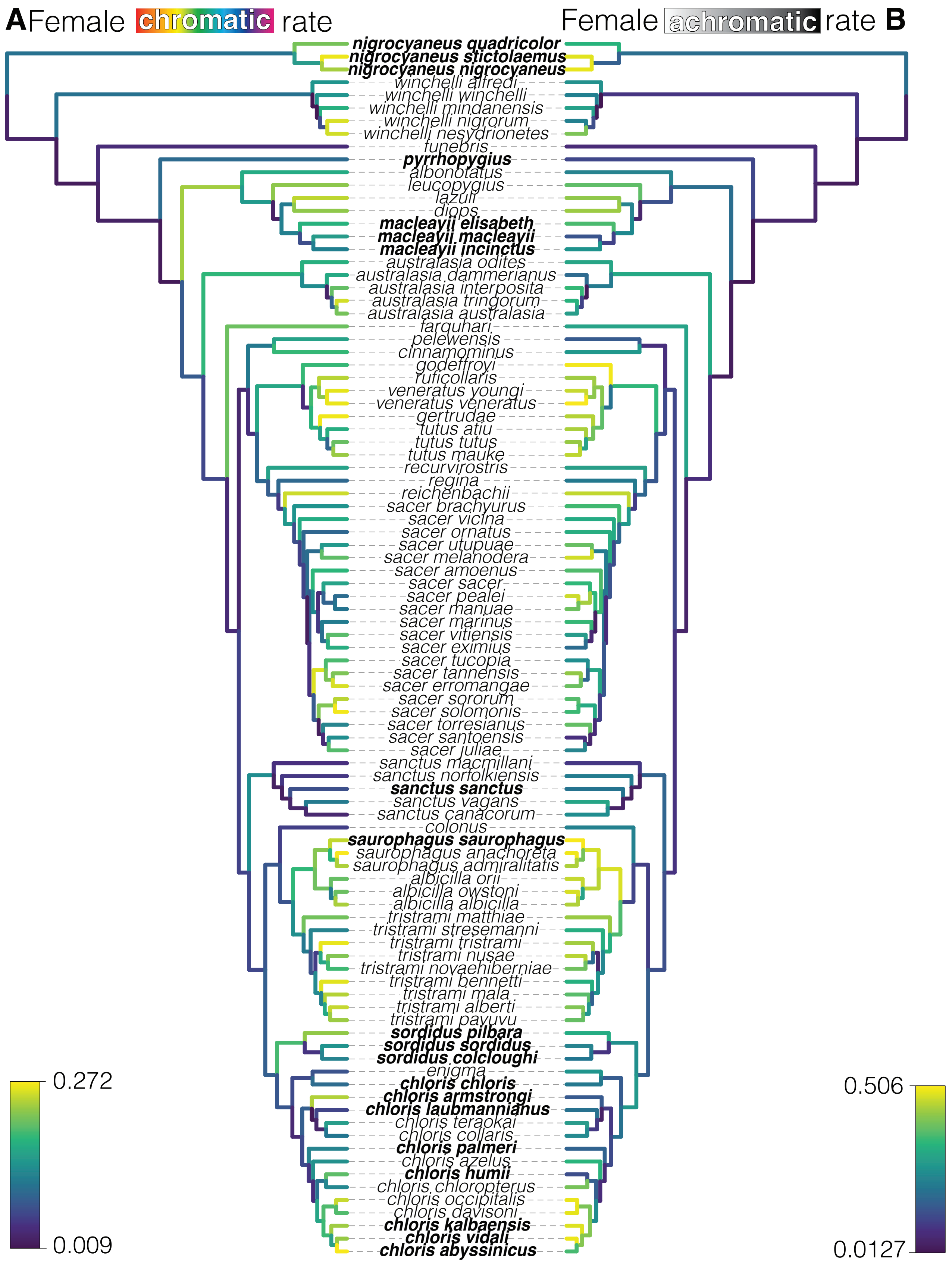


Figure S4. Tempo of female plumage colour evolution for A) chromatic (describing hue and saturation) and B) achromatic (describing brightness) colours across 14 plumage patches. Warmer branch colours indicate higher rates of colour evolution. Bolded names indicate continental taxa.

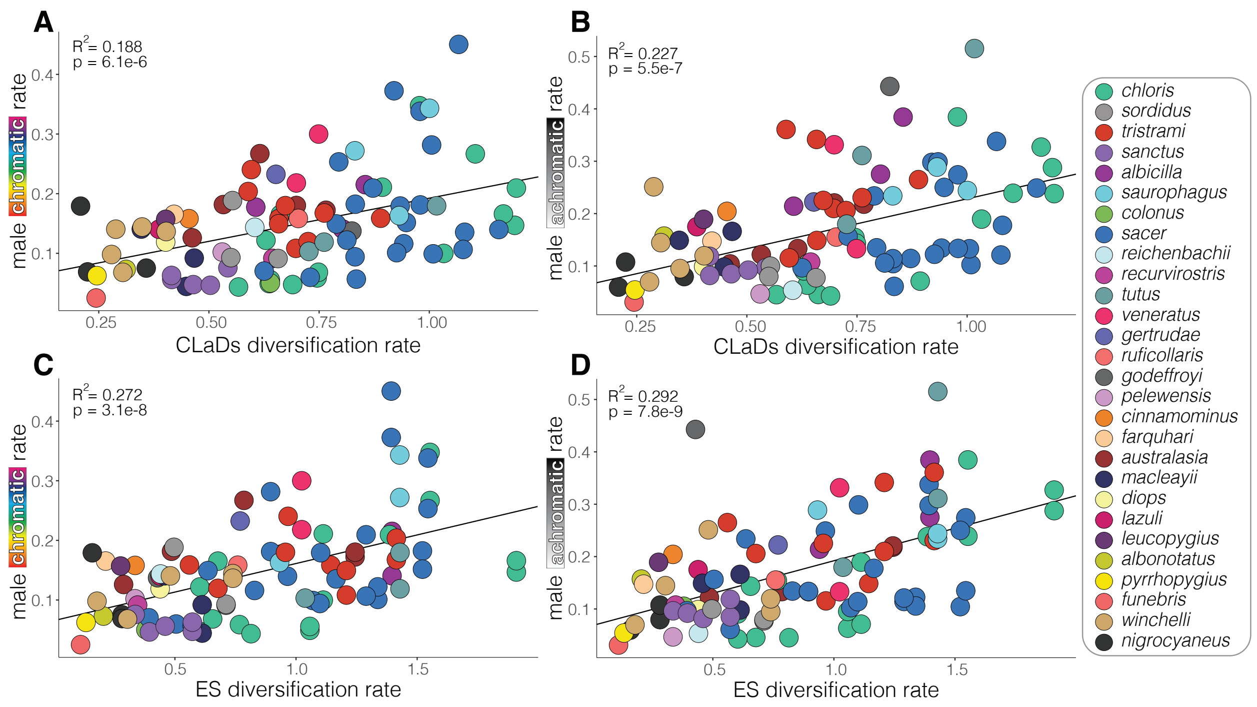


Figure S5. Positive relationship between diversification rate and male chromatic (A,C) and achromatic (B, D) evolutionary rates in Todiramphus. Top panels correspond to lineage diversification rates estimated with cladogenetic diversification rate shift (CLaDs) model and the bottom panels are rates inferred with inverse equal splits (ES). Trend lines, R2, and p-values represent phylogenetic linear model results (See Table S2 for all phylolm results and Table S3 for linear model results). Dot colours correspond to species-level taxonomy. Note panels A and B correspond to those shown in Fig. 1B.

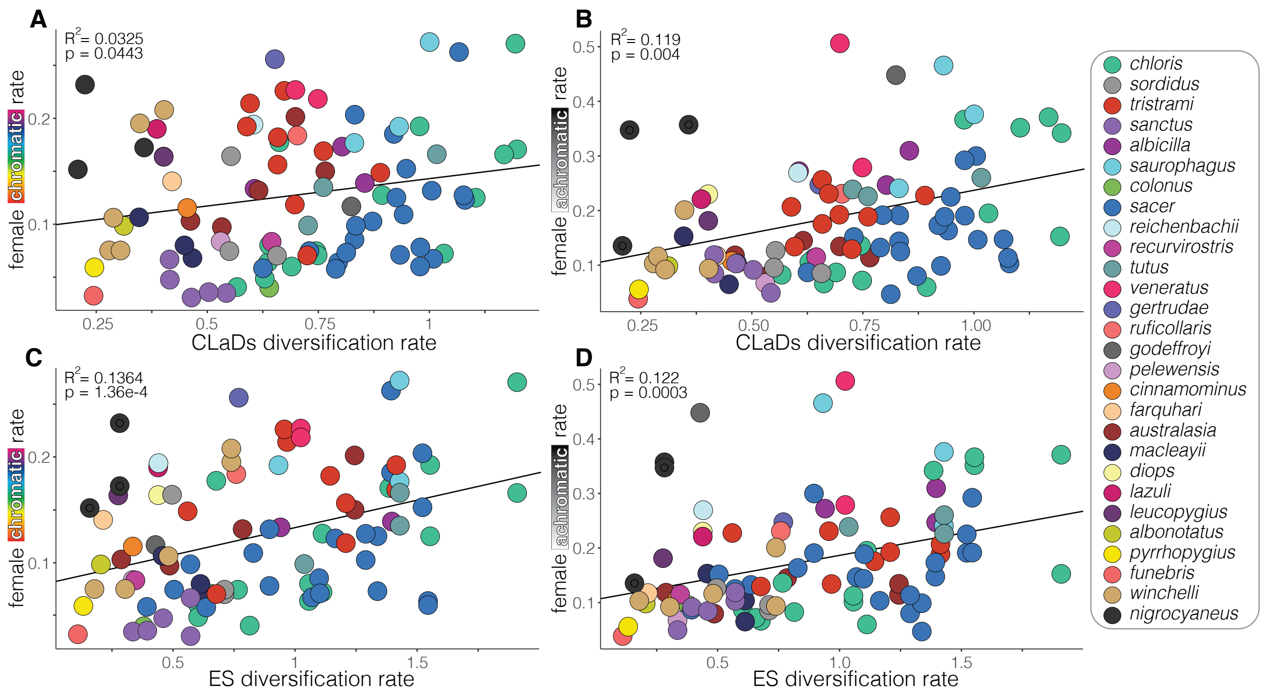


Figure S6. Positive relationship between diversification rate and female chromatic (A,C) and achromatic (B, D) evolutionary rates in Todiramphus. Top panels correspond to lineage diversification rates estimated with cladogenetic diversification rate shift (CLaDs) model and the bottom panels are rates inferred with inverse equal splits (ES). Trend lines, R2, and p-values represent phylogenetic linear model results (See Table S2 for all phylolm results and Table S3 for linear model results). Dot colours correspond to species-level taxonomy.

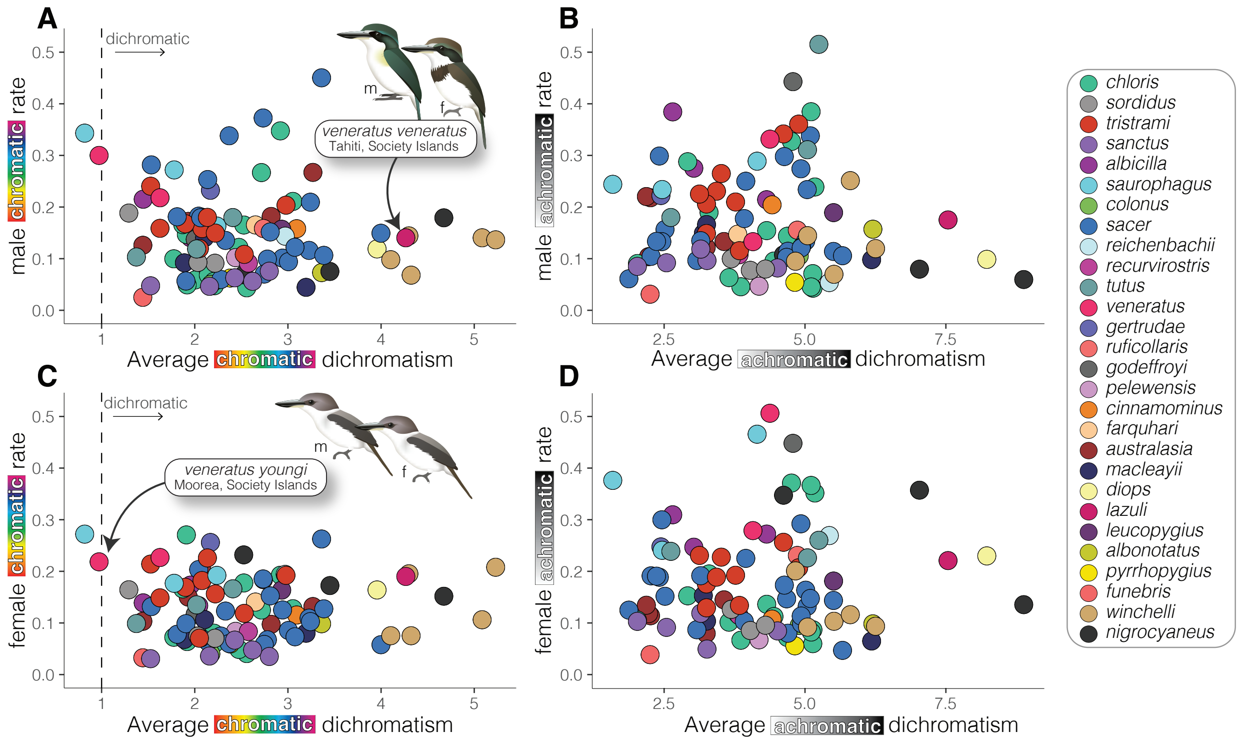


Figure S7. No relationship between the degree of sexual dichromatism and tempo of male (A–B) or female (C–D) plumage evolutionary rates. Chromatic dichromatism (left panels) describes differences in hue and saturation between the sexes whereas achromatic dichromatism (right panels) describes differences in brightness. Dichromatism values larger than one indicate taxon is noticeably different in the avian visual system. Dot colours correspond to species-level taxonomy. Note that all taxa are identifiably dichromatic for achromatic colour. Example taxa show differences in chromatic dichromatism between the two described taxa of Society Kingfisher (*T. veneratus*), in which *T. v. youngi* is one of two its two taxa to not be identifiably dichromatic for hue and saturation. Illustrations created by Jenna McCullough.

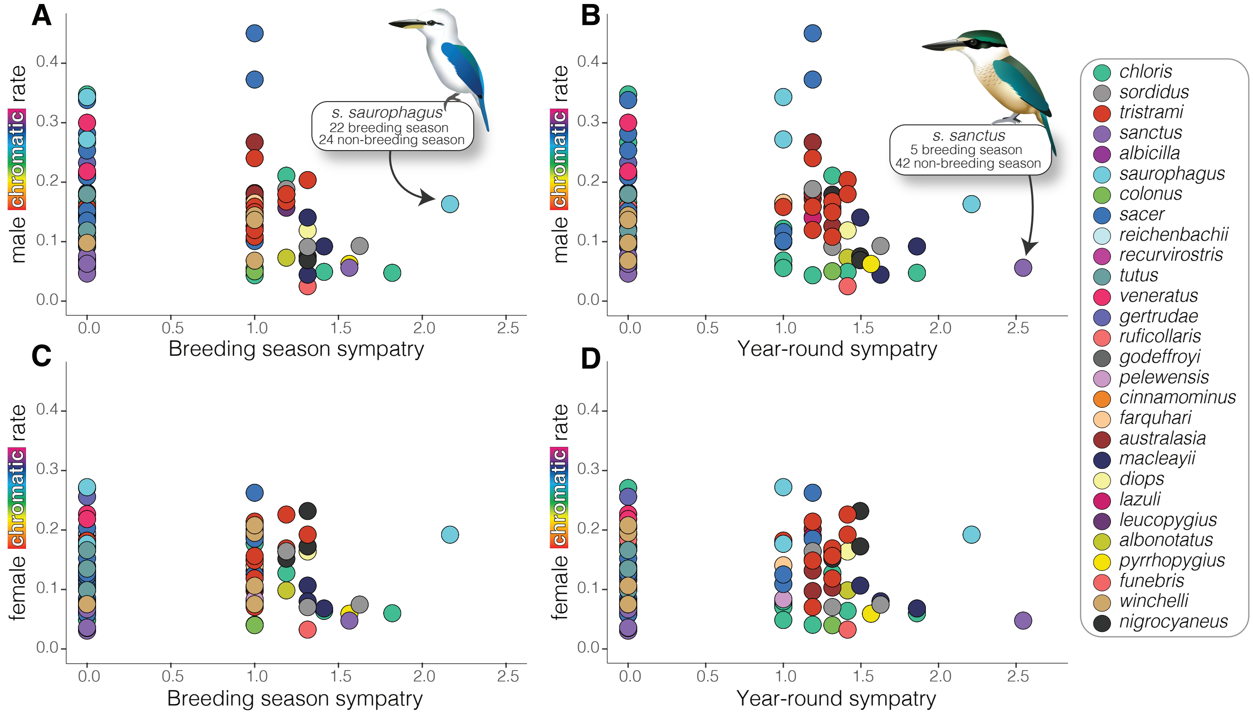


Figure S8. No relationship between male (top panels) or female (bottom) chromatic evolutionary rates and number of sympatric taxa during the breeding (left) and non-breeding (right) season. Dot colours correspond to species-level taxonomy. Sympatry data is fourth-root transformed. Values of zero indicate that the taxon is allopatric and does not come into contact with another Todiramphus species. Illustrations of the widespread and nominate taxon of Beach Kingfisher (*T. saurophagus saurophagus*) and the Sacred Kingfisher (*T. sanctus*), an austral migrant taxon, were created by Jenna McCullough.

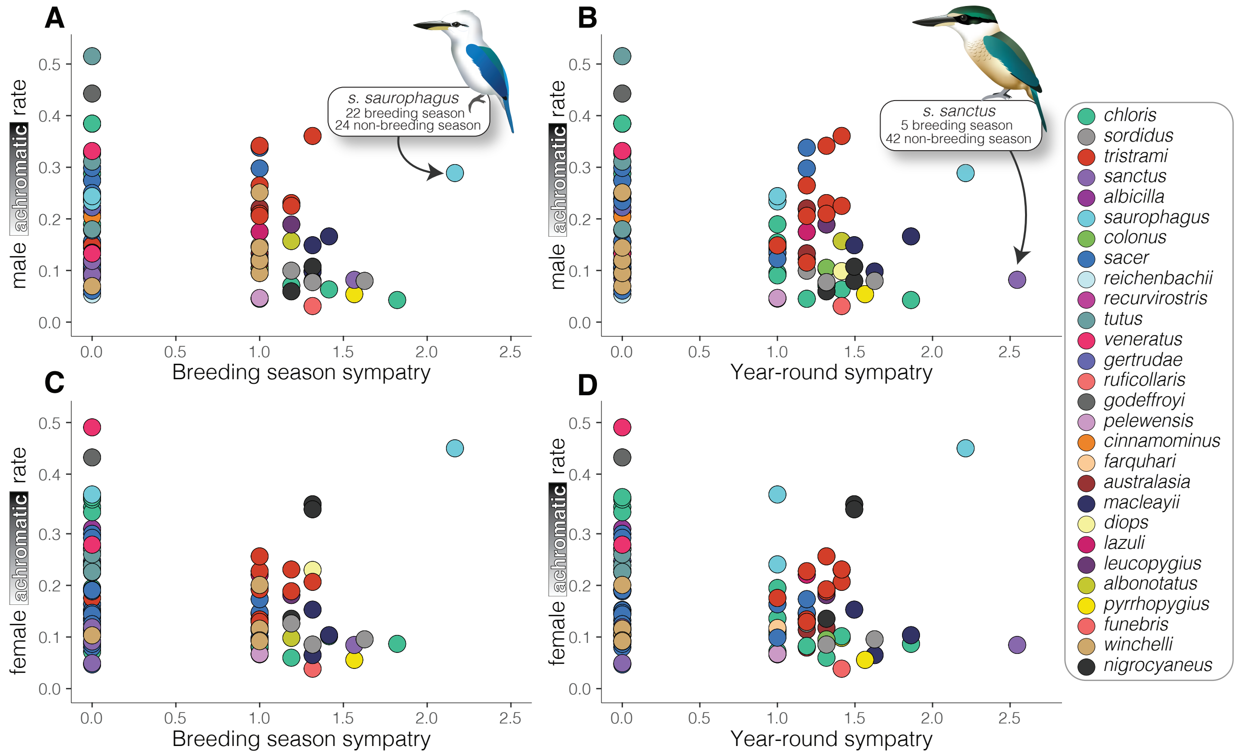


Figure S9. No relationship between male (top panels) or female (bottom) achromatic evolutionary rates and number of sympatric taxa during the breeding (left) and non-breeding (right) season. Dot colours correspond to species-level taxonomy. Sympatry data is fourth-root transformed. Values of zero indicate that the taxon is allopatric and does not come into contact with another Todiramphus species. Illustrations of the widespread and nominate taxon of Beach Kingfisher (*T. saurophagus saurophagus*) and the migratory taxon of Sacred Kingfisher (*T. sanctus*) were created by Jenna McCullough.

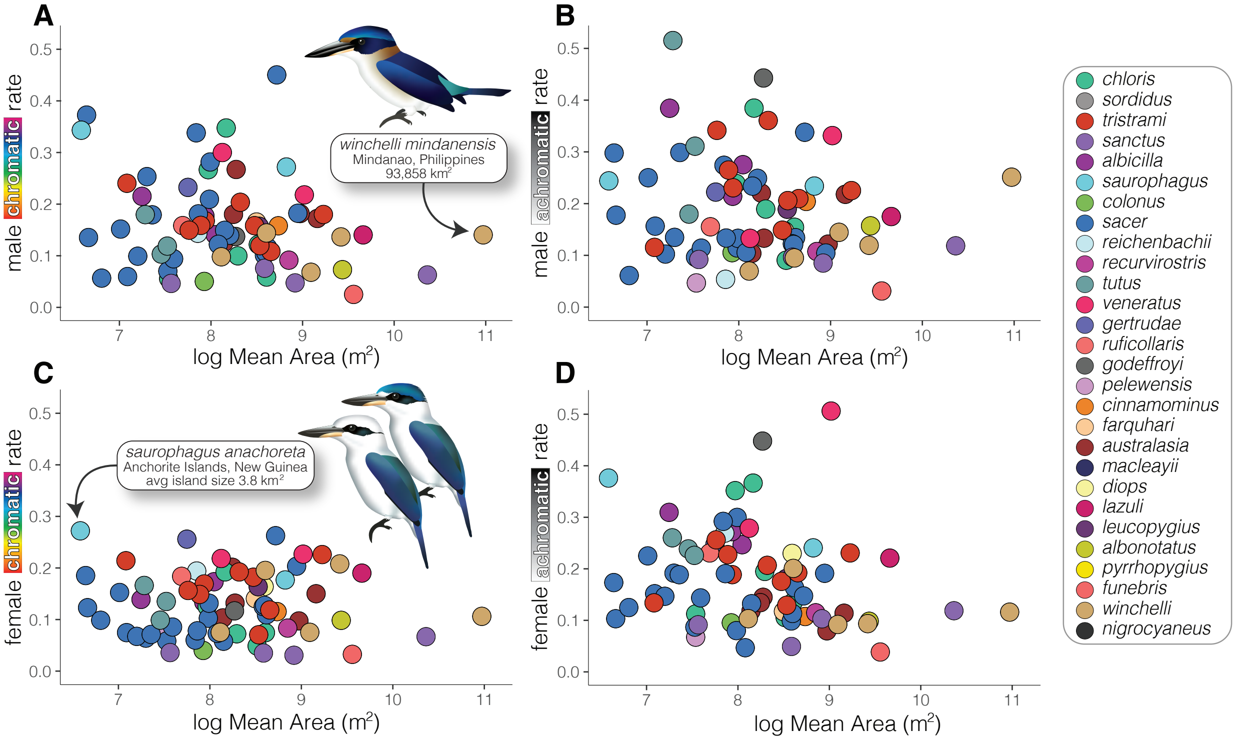


Figure S10. No statistically supported relationship between male (top) or female (bottom) chromatic (left) and achromatic (right) evolutionary rates for log transformed mean island size (m$^2$). Dot colours correspond to species-level taxonomy. Illustrations describe range in mean island size: the Rufous Collared Kingfisher (*T. winchelli mindanensis*) is endemic to only Mindanao, Philippines whereas one taxon of the Beach Kingfisher (*T. saurophagus anachoreta*) is endemic to the Anchorite islands, Papua New Guinea. *T. s. anachoreta* is one of two Todiramphus taxa that have dual plumage morphotypes for crown color (see Supplementary methods), both are shown in panel C. Illustrations created by Jenna McCullough.

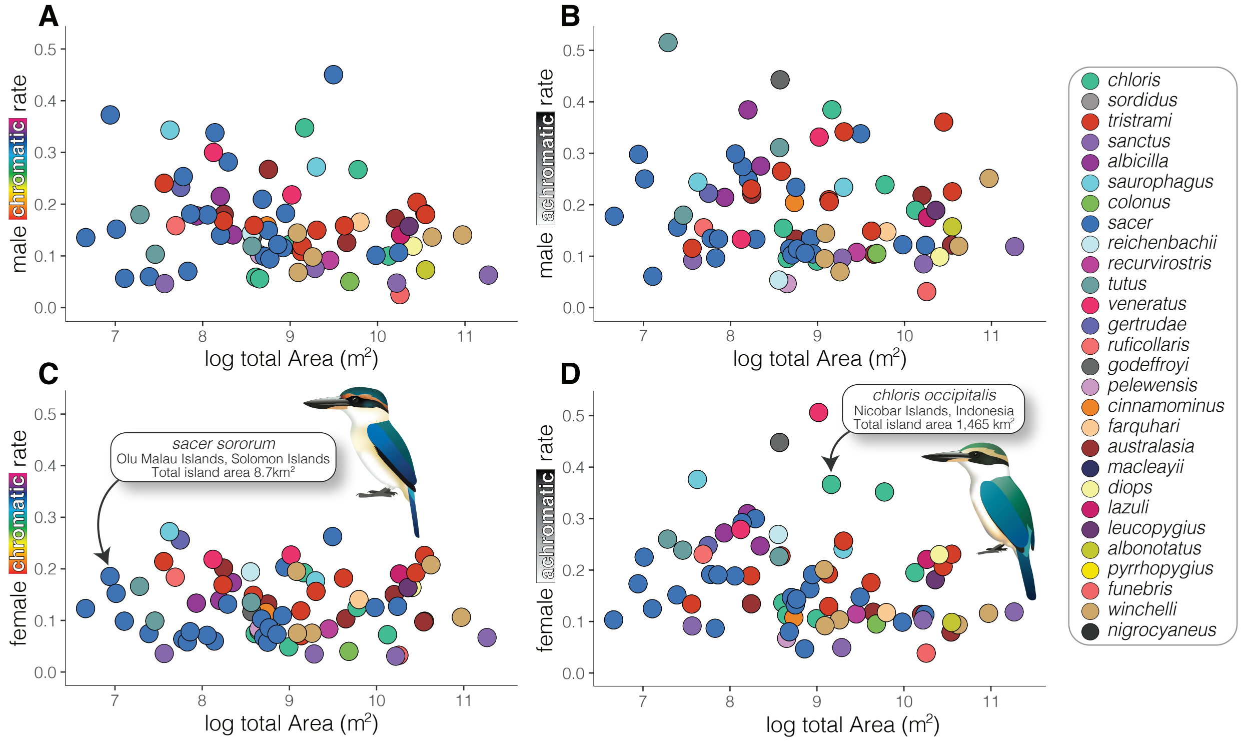


Figure S11. No statistically supported relationship between male (top) or female (bottom) chromatic (left) and achromatic (right) evolutionary rates for log transformed total island size (m$^2$). Dot colours correspond to species-level taxonomy. Illustrations demonstrate a range in total island size: one taxon of Pacific Kingfisher (T. s. sororum) is endemic to the Olu Malau Islands, Solomon Islands and one taxon of Collared Kingfisher (T. chloris occipitalis) is endemic to the Nicobar Islands, Indonesia. Illustrations created by Jenna McCullough.

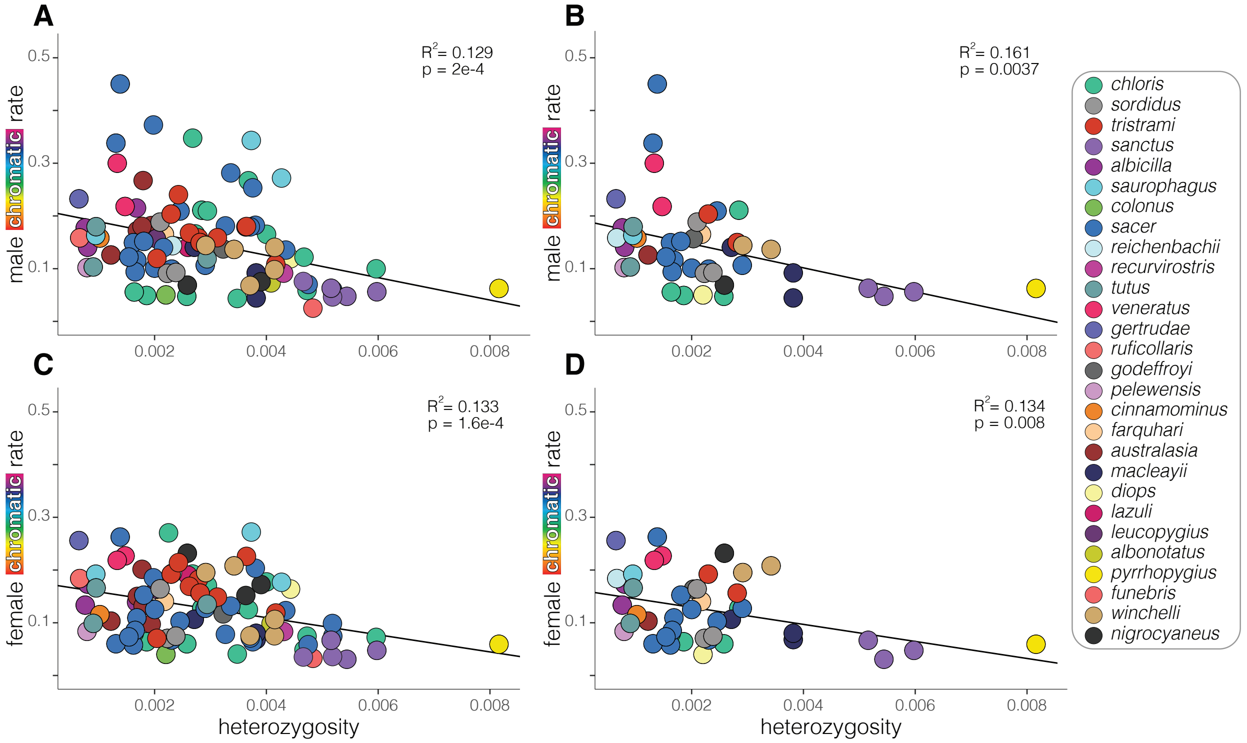


Figure S12. Less genetically diverse populations have higher chromatic (hue, saturation) evolutionary rates. Male (top) and female (bottom) chromatic evolutionary rates with all taxa (left side). When taxa whose heterozygosity estimates are sourced from toepads were removed (right side), the relationship is still statistically significant. Dot colours correspond to species-level taxonomy. See Table S2 for all phylogenetic linear models and Table S3 for all linear models. Note that Panel A corresponds to Fig. 2C.

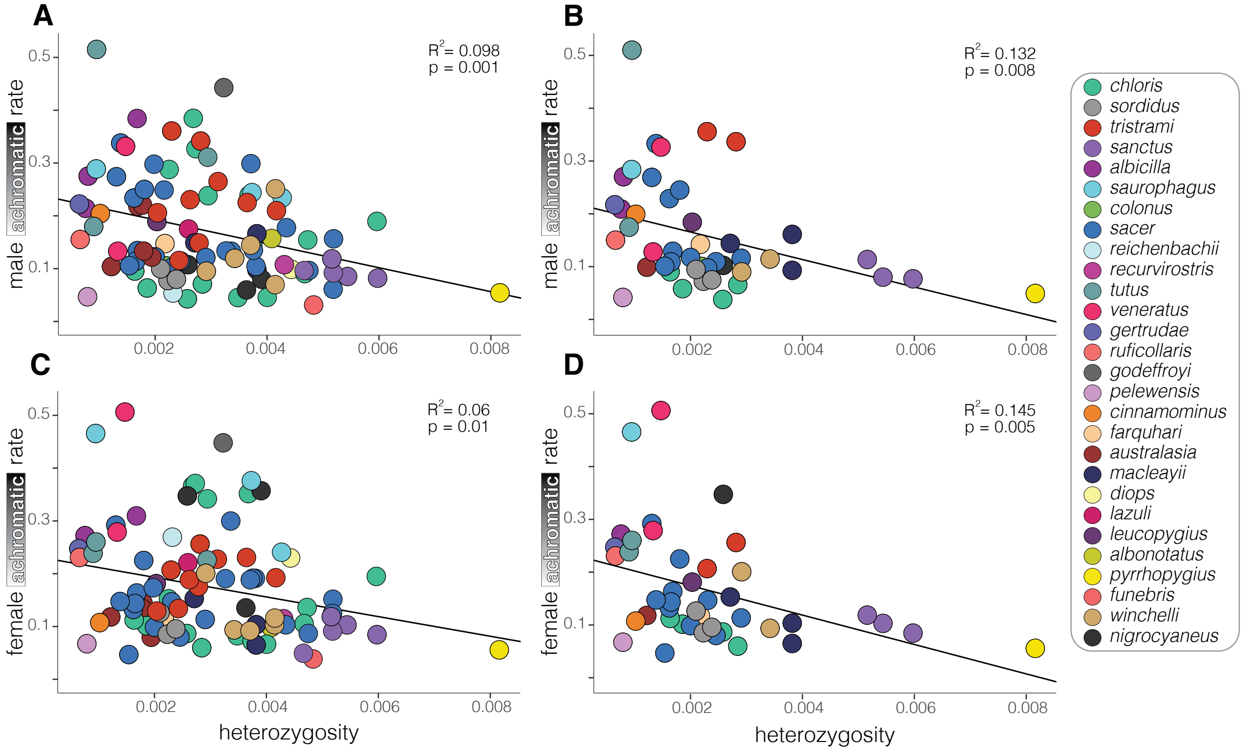


Figure S13. Less genetically diverse populations have higher achromatic (brightness) evolutionary rates. Male (top) and female (bottom) chromatic evolutionary rates with all taxa (left side). When taxa whose heterozygosity estimates are sourced from toepads were removed (right side), the relationship is still statistically significant. Dot colours correspond to species-level taxonomy. See Table S2 for all phylogenetic linear models and Table S3 for all linear models. Note that Panel A corresponds to Fig. 2F.

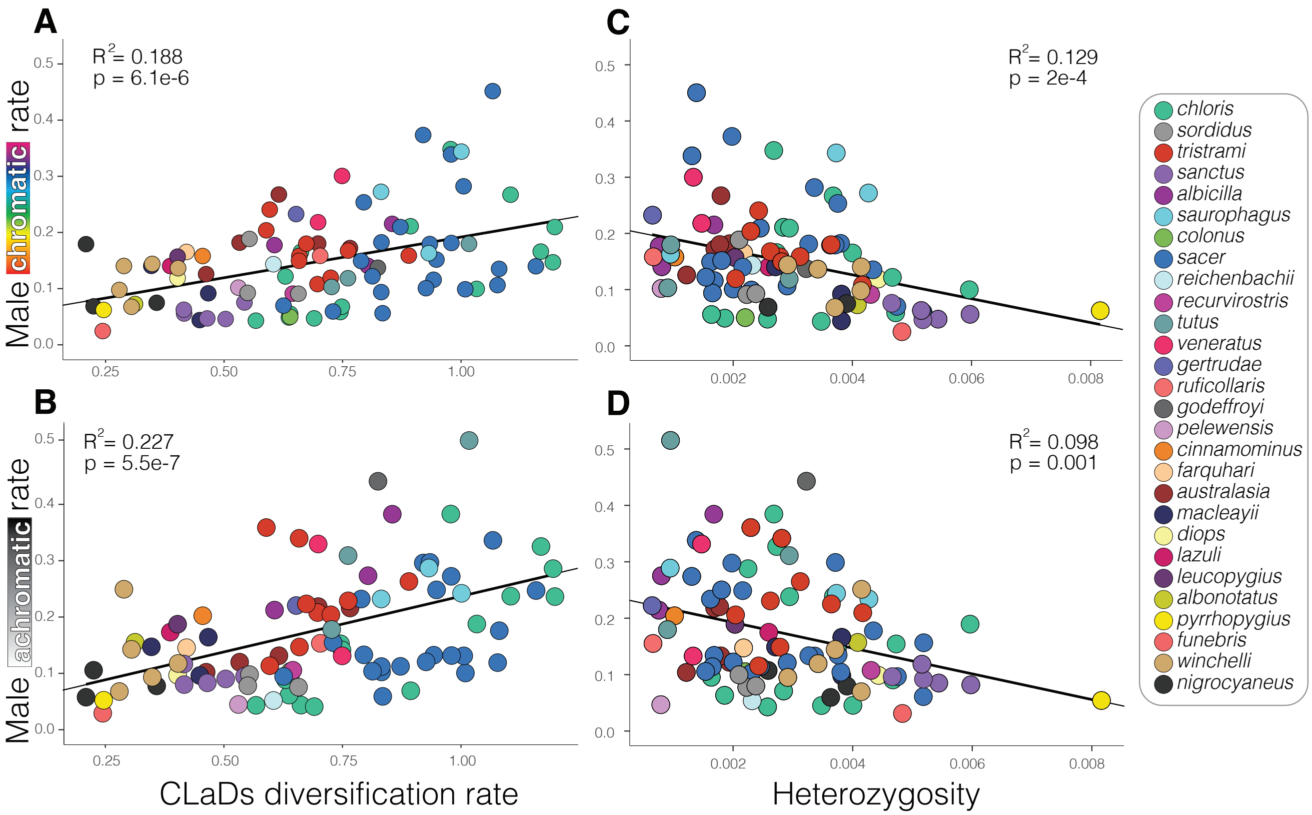


Figure S14. Plumage colour evolutionary rates are higher in more rapidly diversifying lineages that have lower genomic diversity. Male chromatic (A) and achromatic (B) rates are positively correlated with diversification rates and negatively correlated with genome-wide estimates of heterozygosity. Dot colours correspond to species-level taxonomy. Note that Panels A and B correspond to Fig. 1B and Panels C and D correspond to Fig. 2C,F.

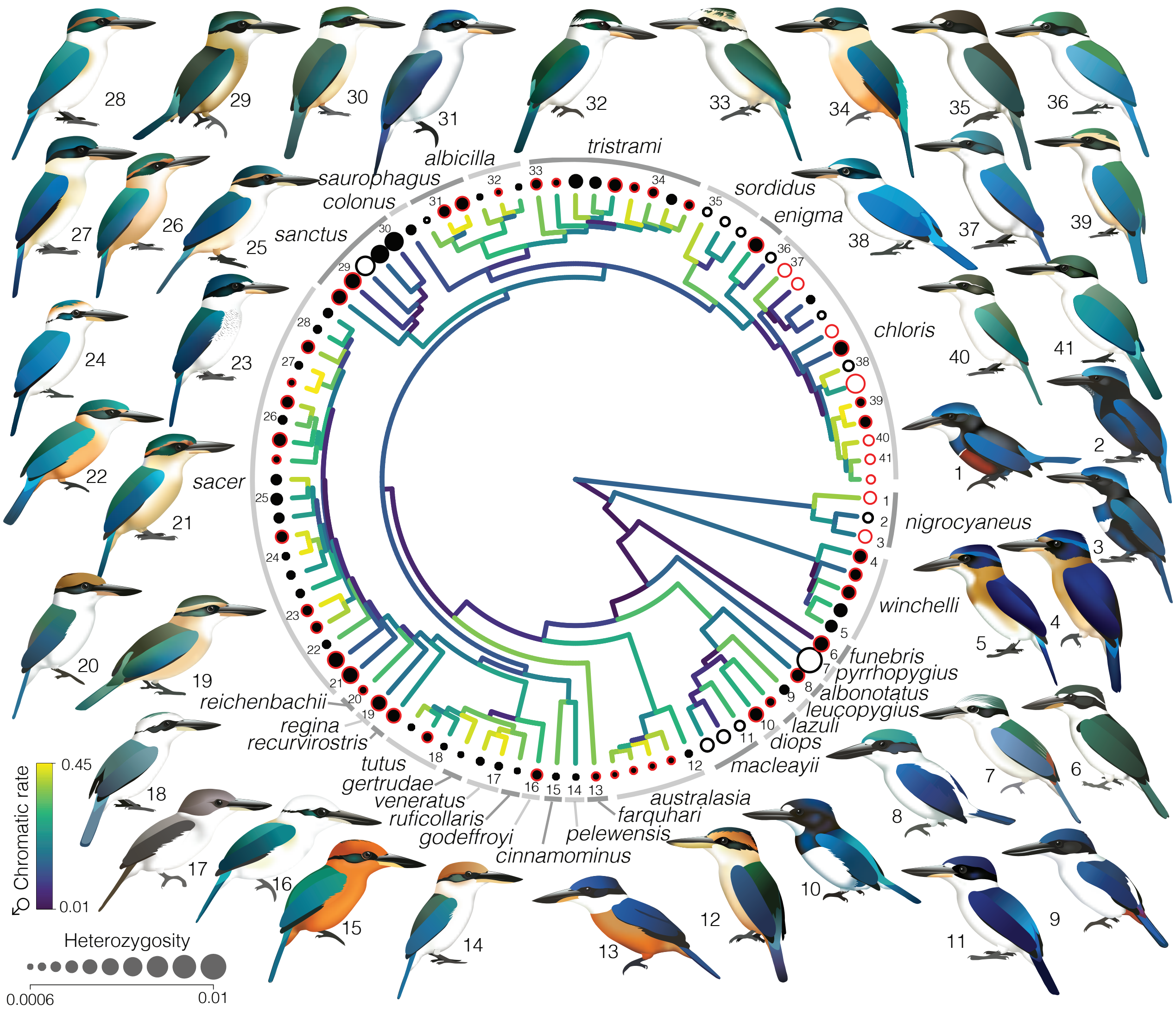


Figure S15. Lineages with smaller population sizes have higher rates of plumage colour evolution. Phylogeny of maximum clade credibility (MCC) timetree, with branch colours showing the ancestral state reconstruction of colourspace XYZ coordinates for males. Warmer branch colours indicate higher rates of plumage colour evolution. Size of circles at tips reflect genome-wide estimates of heterozygosity (Pi). Colour of dots indicate if a species is island endemic (black fill) or occurs on continental landmasses (white). The border of dots indicate the source of genomic data, whether it was estimated from a high-quality frozen tissue sample (black) or cut from toe pads of museum study skins (red). Illustrated *Todiramphus* taxa were created by Jenna McCullough and are as follows: 1) *T. nicrocyanus quadricolor,* 2) *T. nigrocyanus stictolaemus,* 3) *T. nigrocyaneus nigrocyaneus,* 4) *T. winchelli alfredi,* 5) *T. winchelli nesydrionetes,* 6) *T. funebris,* 7) *T. pyrrhopygius,* 8) *T. albonotatus,* 9) *T. leucopygius,* 10*) T. diops,* 11) *T. macleayii elisabeth,* 12) *T. australasia odites,* 13) *T. farquhari,* 14) *T. pelewensis,* 15) *T. cinnamominus,* 16) *T. godeffroyi,* 17) *T. veneratus youngi,* 18) *T. tutus atiu,* 19) *T. regina,* 20) *T. reichenbachii,* 21) *T. sacer brachyurus,* 22) *T. sacer ornatus,* 23) *T. sacer melanodera,* 24) *T. sacer pealei,* 25) *T. sacer vitiensis,* 26) *T. sacer tannensis,* 27) *T. sacer solomonis,* 28) *T. sacer santoensis,* 29) *T. sanctus norfolkiensis,* 30) *T. sanctus canacorum,* 31) *T. saurophagus anachoreta,* 32) *T. albicilla owstoni,* 33) *T. tristrami matthiae,* 34) *T. tristrami mala,* 35) *T. sordidus pilbara,* 36) *T. chloris chloris,* 37) *T. chloris armstrongi,* 38) *T. chloris humii,* 39) *T. chloris occipitalis,* 40) *T. chloris kalbaensis,* 41) *T. chloris vidali.*

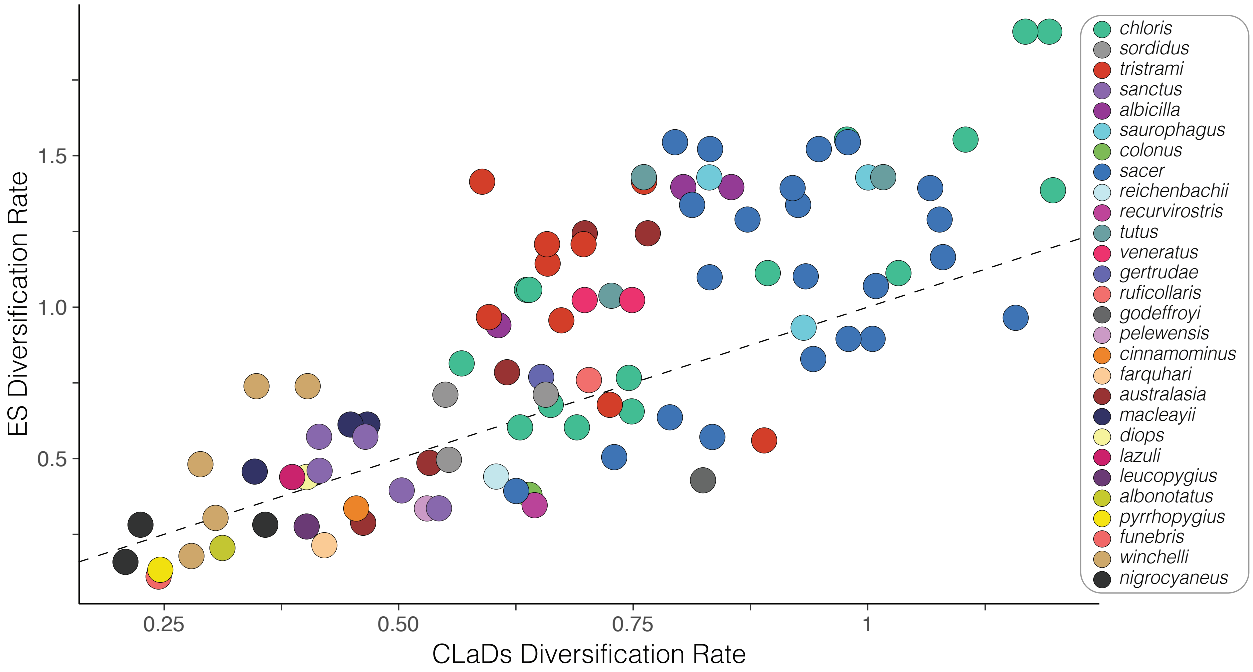


Figure S16. Comparison of diversification rates as inferred by inverse equal splits (ES) and cladogenetic diversification rate shift (CLaDs) models. Dotted line is the line of equivalence. Dot colours correspond to species limits.
